# A genome-wide CRISPR base-editing resource for functional genomics and drug-response mapping in *Leishmania mexicana*

**DOI:** 10.1101/2025.06.16.659792

**Authors:** Jorge Arias del Angel, Fabian Link, Nicole Herrmann May, Ilkem Ekici, Konstantin Wawra, Simon Schwind, Simon Zorn, Jennifer Haggarty, Stefan K Weidt, Ryan Ritchie, Michael P Barrett, Ger van Zandbergen, Tom Beneke

## Abstract

We present a CRISPR/Cas9 cytosine base editing library for genome-wide loss-of-function screening in *L. mexicana* and apply it to dissect the genetics of *Leishmania* drug response. The results, accessible at https://www.LeishBASEeditDB.net, revealed novel response biomarkers for Sb^III^, miltefosine, amphotericin B, pentamidine, and the arylmethylaminosteroid 1c. We identified hundreds of loci linked to altered drug responses, including opposing effects among paralogs, cross-resistance, and collateral sensitivity. Among 41 validated candidates, we identified sterol defects in two novel amphotericin B markers, discovered a regulator of intracellular miltefosine transporter complex localization, and uncovered evidence that flagellar-associated defects reduce drug sensitivity. Parallel viability and motility screens revealed, for the first time, genome-wide fitness contributions of unique and multi-copy genes in *Leishmania* promastigotes. Our approach enables powerful reverse genetic screens across *Leishmania* species, advancing drug mechanism studies and guiding combination therapy designs. The library is available for others to screen a multitude of additional loss-of-function phenotypes.

## INTRODUCTION

Many parasitic diseases face increasing challenges from emerging drug treatment failure, and *Leishmania* spp., protozoan parasites transmitted by infected sand flies, represent an excellent experimental model for studying antiparasitic resistance due to their clinical relevance and the diverse mechanisms underlying treatment failure (*1–8*). In *Leishmania*, resistance to key drugs, including antimonials, pentamidine, miltefosine, and amphotericin B, frequently arises through alterations in drug influx and efflux systems, notably involving ATP-binding cassette (ABC) transporters (*9–13*), the aquaglyceroporin AQP1 (*14–23*) and the miltefosine transporter complex (LMT) (*24–28*). Resistance can also associate with changes in membrane composition, particularly lipid and sterol profiles in the case of amphotericin B and miltefosine (*29–41*).

Beyond membrane dynamics, resistance in *Leishmania* can be transferred via extracellular vesicles (*42, 43*) and often involves gene copy number variations or point mutations in genes associated with transcription, translation, mitochondrial activity, metabolic networks, and cell signalling (*2, 44–51*). These mechanisms are largely drug-specific, though some mutations have been linked to cross-resistances (*13, 39, 52–55*) (e.g. in AQP1, LMT, and various ABC transporters).

By contrast, few studies have examined mutations that increase drug sensitivity, and whether resistance to one drug routinely alters susceptibility to others remains unclear, highlighting a gap in understanding collateral sensitivities (*17, 38, 56, 57*). Importantly, defining the genetic basis of both resistance and sensitivity can guide treatment strategies, including combination therapies (*1, 8, 38, 54, 57*).

To address this knowledge gap, we conducted a genome-wide functional analysis in *L. mexicana* using our cytosine base editing (CBE) platform, overcoming limitations inherent in *Leishmania* that preclude standard CRISPR or the RNAi approaches (*58, 59*) that revolutionised understanding of drug-response mechanisms in the related parasite *Trypanosoma brucei* (*60*). Our screen identified dozens of novel resistance and hypersensitivity biomarkers in *L. mexicana*, while parallel viability screening provided unprecedented genome-wide mapping of fitness-related genes. The transfected parasite library can be easily revived and represents a resource available for future studies.

## RESULTS

### A GENOME-WIDE LOSS-OF-FUNCTION SCREEN FOR PHENOTYPING IN *LEISHMANIA*

To enable genome-wide loss-of-function screening in *L. mexicana*, we generated a library comprising 38,422 protein-targeting CBE sgRNAs (supplementary file 1), with 1 to 6 guides per gene (∼5.2 average). These sgRNAs introduce STOP codons within the first 50% of coding sequences in 94% of all protein-coding ORFs, alongside 1,000 non-targeting controls. The library was integrated into the 18S rRNA SSU locus via Cas12a-mediated recombination at 500 parasites per sgRNA. With one sgRNA expressed per cell (*59*), functional assessment of 7,718 gene targets was enabled (fig. 1A-B; supplementary file 2; data available at https://www.LeishBASEeditDB.net).

**Figure 1.**
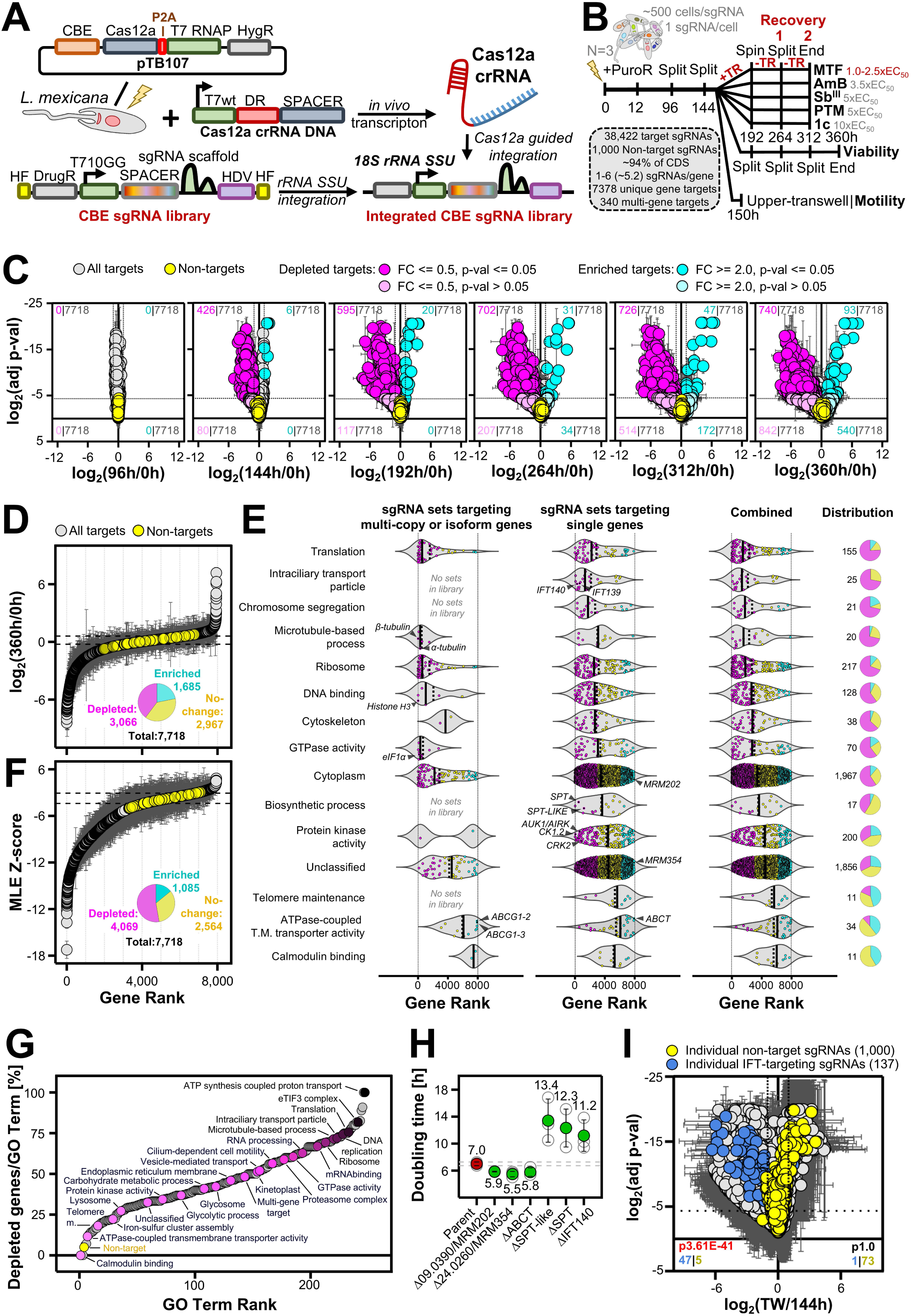
Base editing enables genome-wide loss-of-function screening in *L. mexicana*. **(A)** Schematic representation of Cas12a-mediated integration of the CBE sgRNA expression library, as described previously (*59*). **(B)** Workflow of the genome-wide loss-of-function screen performed in *L. mexicana* across three independent biological replicates. “+TR” and “−TR” indicate time points with and without drug treatment, respectively, for each anti-leishmanial screen. **(C)** Median values of sgRNA sets targeting 7,718 unique or multi-gene targets shown. Error bars represent standard deviations from three biological replicates. The number of enriched or depleted sgRNA sets is indicated for each panel, using fold-change thresholds of ≥ 2 and ≤ 0.5, respectively. Statistically significant differences were defined by p-values ≤ 0.05, while those with higher p-values were considered a secondary enriched or depleted category. Comparisons include plasmid reference (0 h) vs. post-transfection time points (96 h, 144 h, 192 h, 264 h, 312 h, 360 h). **(D)** Median sgRNA set log₂ fold-change values plotted against ranked positions (0 h vs. 360 h post-transfection). Gray: 7,718 unique or multi-gene targets; yellow: non-target controls; error bars: standard deviations from three biological replicates; dotted line: 95% reference interval based on non-target log₂ fold-changes. The pie chart shows the percentage of sgRNA sets below (depleted), above (enriched), or within (no-change) the reference interval. **(E)** Dots within violin plots indicate ranked positions of sgRNA sets from **(D)** for each GO term. Depleted sets are shown in magenta, enriched in cyan, and others in yellow; dotted lines: mean; solid lines: median. The far-right column shows total genes screened and their distribution across fitness categories. **(F)** Median sgRNA set MLE z-scores plotted against ranked positions; all as in **(D)**. **(G)** The percentage of depleted genes per GO term class (with a minimum of 10 genes/class) is plotted against its average rank across replicates. **(H)** Doubling times of generated null mutants. Error bars represent standard deviation across triplicates. **(I)** Comparison of the 144 h post-transfection sample vs. the upper transwell fraction. Highlighted are non-targeting controls and sgRNA sets associated with the GO term “Intraciliary transport particles” (GO:0030990), with the number of significantly enriched or depleted sgRNAs indicated (using thresholds of ≤−1 or ≥1 for log_2_ fold change and ≤−4.3219 for log_2_ p-value). P-value from two-sided Fisher’s exact tests (2×2 contingency tables) are shown for each group.

We evaluated the functionality of the library by assessing promastigote fitness through serial sub-culturing over 360 h (fig. 1B). Initial quality control at the individual sgRNA level confirmed high mapping rates, low Gini indices, and even sgRNA-read distributions across samples and replicates (fig. S1A-E, S2A; supplementary file 2). Pairwise (PW) assessment of sgRNA-level enrichments showed stable behaviour of non-targeting controls over time (fig. S3A; supplementary file 3), supporting subsequent aggregation of sgRNAs into gene-targeting sets (7,378 unique and 340 multi-copy or isoform genes) by calculating gene-level log_2_ fold changes as the median of individual sgRNA values (fig. 1C; supplementary file 4). Ranking gene-level effects from the 360 h versus 0 h PW comparison identified 39.7% (3,066) of targets as depleted and 21.8% (1,685) as enriched over non-targeting controls (fig. 1D; supplementary file 5A).

To confirm the biological relevance of these phenotypes, we ranked to assess the enrichment of Gene Ontology (GO) terms associated with essential functions. Genes linked to processes such as “Translation”, “Intraciliary Transport Particle” (IFT), “Chromosome segregation”, “Microtubule-based process”, “Ribosome”, “DNA binding”, and “Cytoskeleton” were enriched among depleted hits, consistent with essential roles (fig. 1E&G; supplementary file 6). These included well-characterized fitness-associated genes like IFT140 (LmxM.31.0310), IFT139 (LmxM.04.0550), CRK2 (LmxM.05.0550), AUK1/AIRK (LmxM.28.0520), and CK1.2 (LmxM.34.1010) (fig. 1E) (*59, 61–67*). Our library also effectively targeted multi-copy or isoform genes using overlapping sgRNA sets. Alpha-(LmxM.13.0280-0300) and beta-tubulin (LmxM.08.1171, LmxM.08.1230, LmxM.21.1860, LmxM.32.0792 & LmxM.32.0794), histone H3 (LmxM.10.0870, LmxM.10.0970, LmxM.10.0990 & LmxM.16.0570-0610), and eIF1α (LmxM.17.0080-0086), all essential and present in dispersed copies or tandem repeats, were significantly depleted (fig. 1E) (*68–71*). In contrast, targeting ABCG1-3 isoforms (LmxM.06.0080-0100) or ABCG1&2 alone resulted in a gain-of-fitness phenotype (fig. 1E). GO analysis further indicated that targeting genes associated with “telomere maintenance” and “calmodulin binding”, processes that may be mechanistically-linked (*72, 73*), consistently increased fitness, suggesting these pathways constrain promastigote proliferation under our assay conditions (fig. 1E&G).

While the 360 h versus 0 h PW comparison captured depletion and enrichment effectively, we also assessed temporal dynamics of sgRNA abundance. Using MAGeCK MLE (*74*), we calculated median z-scores of gene-targeting sgRNA sets across all time points (fig. 1C) and compared them to the non-targeting control interval, revealing consistent temporal patterns (fig. 1F; supplementary file 5B). This approach identified a larger fraction of depleted genes (52.7%; 4,069) but fewer enriched genes (14.1%; 1,085).

To validate findings, we individually deleted six genes using Cas12a-mediated gene replacement (fig. S4). Doubling times of mutants (fig. 1H) correlated with screen results (fig. 1E). Given IFT proteins are essential in flagellar assembly (*64–67*), we tested whether motility phenotypes could be resolved. Placing the library into the bottom chamber of a transwell for 6 h (fig. 1B), demonstrated strong depletion of IFT-targeting sgRNAs from the upper chamber, while non-targeting controls became enriched (fig. 1I; supplementary file 3; 47 out of 137 IFT-targeting sgRNAs depleted), clearly distinguishing motile and non-motile cells.

Overall, our library detects diverse phenotypes without requiring additional selective pressure, representing a key advantage over previous pooled loss-of-function high-throughput screens in *Leishmania* (*36*).

### A STRATEGY TO MAP DRUG RESISTANCE AND HYPERSENSITIVITY *LEISHMANIA* MARKERS

Leveraging this, we designed a promastigote screening strategy to identify genes linked to both drug resistance and hypersensitivity across five anti-leishmanials: potassium antimony(III) tartrate (Sb^III^, active against promastigotes without host-dependent reduction), amphotericin B (AmB), miltefosine (MTF), and pentamidine isethionate (PTM), as well as the experimental arylmethylaminosteroid 1c (*75*) (fig. 1B). Drug concentrations were optimized to preserve library diversity, avoid sgRNA bottlenecks, and maintain even sgRNA-read distributions with stable non-targeting controls following 48 hours of treatment and two rounds of recovery without drugs. For MTF, testing 1×, 1.5×, and 2.5× EC_50_, showed that 1× EC_50_ best preserved library diversity (fig. 1B, S1F, S2B, S3B, S5), while optimal concentrations for other compounds were 5× EC_50_ for Sb^III^, 10× EC_50_ for 1c, 3.5× EC_50_ for AmB, and 5× EC_50_ for PTM (fig. 1B, S1G, S2B, S3C-F, S5). Importantly, the transfected promastigote library could be cryopreserved and revived without substantial loss of diversity (fig. S1F, S2C), despite expected depletion of severe fitness mutants, demonstrating its utility as a robust and distributable resource for *Leishmania* research.

Our drug-response screen analysis, identified significantly (p ≤ 0.05) enriched (resistant; fold change ≥ 2) and depleted (hypersensitive; fold change ≤ 0.5) genes by comparing recovery fractions to their viability screen counterparts (192 h vs. recovery 1 and 264 h vs. recovery 2 in fig. 2 and supplementary file 4; combined rank score in fig. S6 and supplementary file 7). We thereby matched samples by time rather than by cell division number, as we calculated relative enrichment within each library sample and normalized across conditions, making it largely independent of differences in population doubling.

**Figure 2.**
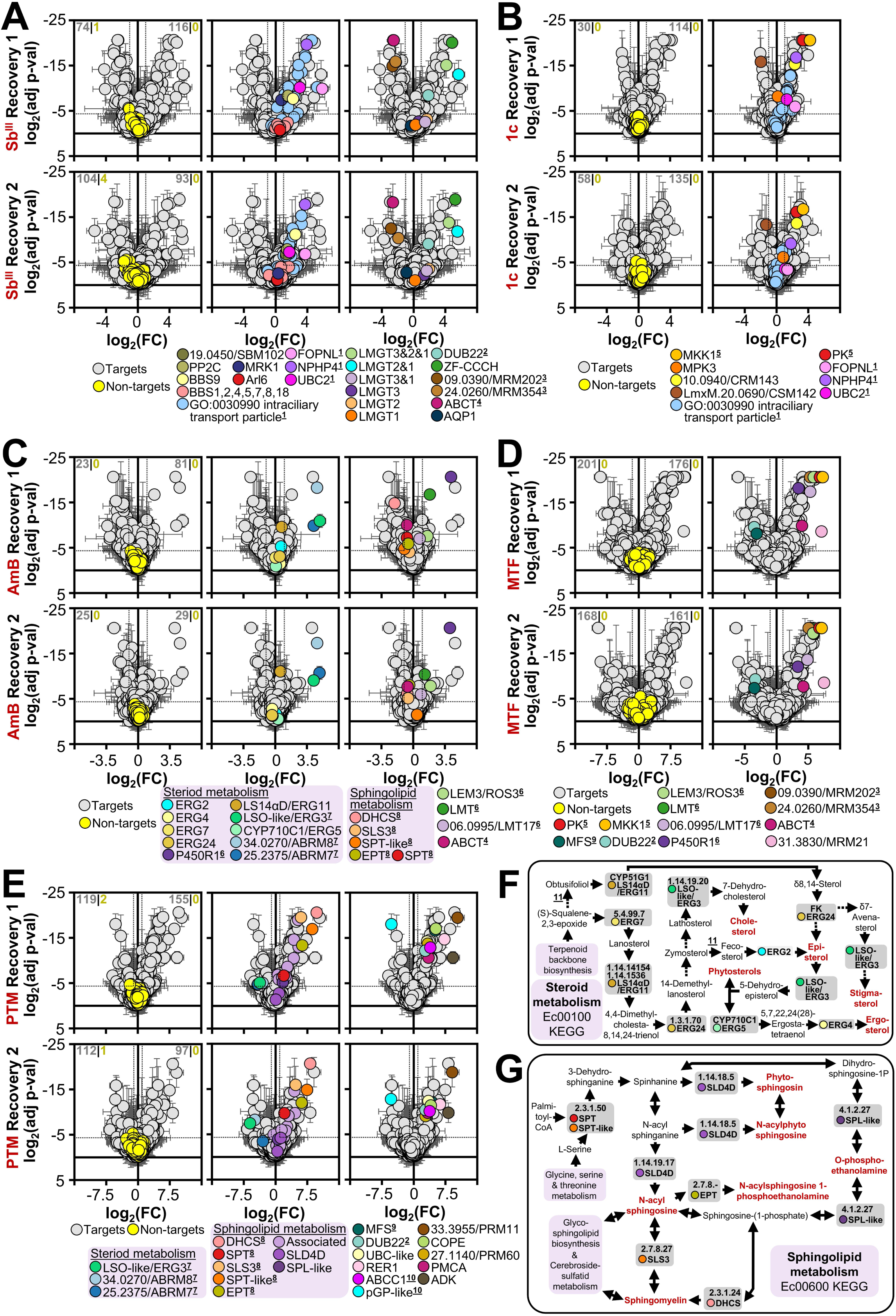
Genome-wide screen identifies *L. mexicana* resistance and hypersensitivity markers. **(A-E)** Median values of sgRNA sets targeting 7,718 unique or multi-gene targets are shown. Error bars represent standard deviations from three biological replicates. The comparison shown is between recovery fractions and their corresponding viability screen time points (192 h vs. recovery 1, 264 h vs. recovery 2). For **(D)**, MTF screen results at 1.0× EC_50_ are shown. Legends below each plot highlight selected genes discussed in the text. The number of significantly enriched or depleted non-targeting (yellow) and targeting (grey) sgRNA sets are indicated in the far left panel of each figure element, using thresholds of ≤−1 or ≥1 for log_2_ fold change and ≤−4.3219 for log_2_ p-value. **(F, G)** Simplified KEGG pathway diagrams for **(F)** steroid metabolism and **(G)** sphingolipid metabolism. Genes from these pathways included in the screen are labelled with their EC numbers and highlighted in grey; matching colours are used for corresponding gene markers in panels **(C)** and **(E)**. Key metabolic products are highlighted in red. **Footnotes: ^1^** Sb^III^ & 1c enriched; **^2^** Sb^III^ & PTM enriched & MTF depleted; **^3^** Sb^III^ depleted & MTF enriched; **^4^** Sb^III^ & AmB depleted & MTF enriched; **^5^**1c & MTF enriched; **^6^** AmB & MTF enriched; **^7^** AmB enriched & PTM depleted; **^8^** AmB depleted & PTM enriched; **^9^** MTF depleted & PTM enriched; **^10^** Paralogous genes with opposing drug response; **^11^** ERG6 (LmxM.36.2380 & 36.2390) not included in screen (no suitable sgRNA found within the 50% of the ORF).

### FLAGELLAR INTEGRITY AND FORMATION DEFECTS CONFER RESISTANCE TO SB^III^ AND 1C

For Sb^III^, we identified multiple genes required for flagellar formation, particularly those associated with intraciliary transport (fig. 1E&I), as strongly linked to resistance (fig. 2A, S6A), reinforcing previous indications that STOP codon insertion in selected IFT genes contributes to resistance (*76*). In addition, the sgRNA-set targeting the IFT regulator BBS9 (LmxM.31.1240) was significantly enriched in both recoveries (fig. 2A), with lower enrichment also observed for BBS2 (LmxM.29.0590), BBS5 (LmxM.12.0680), and BBS7 (LmxM.08_29.0710) (fig. S6A). We then individually deleted all core BBSome components (fig. S4) and confirmed that BBS9 and the slow-growing IFT140 (LmxM.31.0310) mutants (fig. 1H) conferred resistance, whereas complemented knockouts (addbacks [AB]) and other BBSome deletions (BBSome (BBS1 [LmxM.34.4180], BBS2, BBS4 [LmxM.36.2280], BBS5, BBS7, BBS8 [LmxM.09.0440], BBS18 [LmxM.14.0200] (*77*)) did not (fig. 3A). These findings align with studies implicating IFTs and the BBSome in drug resistance in *C. elegans* (*78, 79*) and support an essential role of BBS9 in BBSome assembly and function (*77, 80*) in *Leishmania*. Given that IFT140 (LmxM.31.0310) deletion disrupts *Leishmania* flagellar pocket morphology (*66*), the site of all endo- and exocytosis (*81, 82*), we next tested whether genes involved in pocket function, including flagellar attachment zone (FAZ) components (*83*) and vesicle trafficking factors (e.g., SNAREs, Rabs, VPS) (*84, 85*) conferred resistance. However, other than minor effects with Rab5a (LmxM.18.1130) and Rab6 (LmxM.02.0260) (fig. S7) no impact was seen.

**Figure 3.**
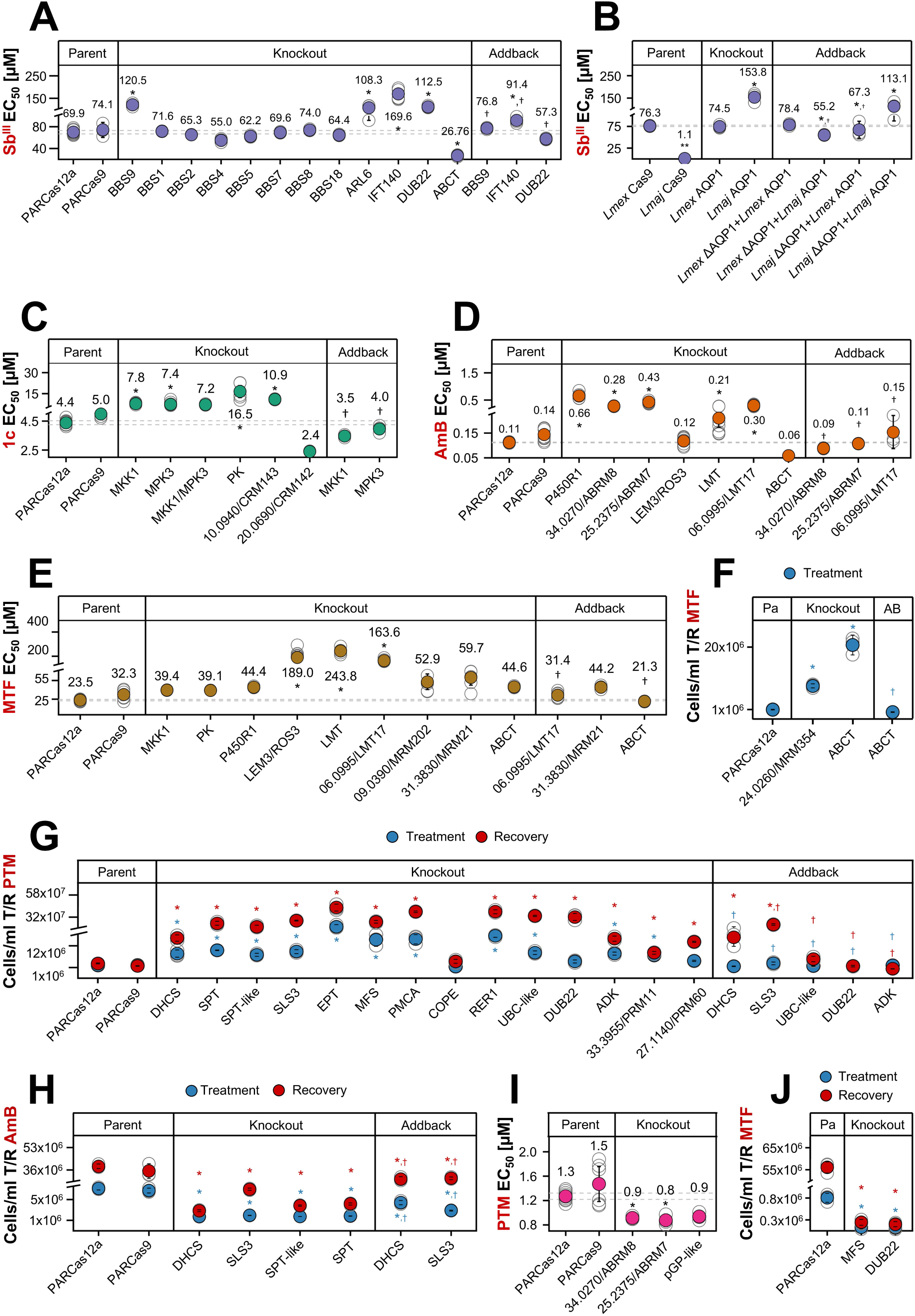
Individual mutant verification confirms data from genome-wide screen. **(A-E, I)** EC_50_ values are shown for each tested cell line following an MTT assay, including parental controls, knockouts, and addbacks as indicated. Each filled-colored dot represents the mean EC50 from up to nine biological replicates (individual replicates shown as grey unfilled dots), with error bars representing the 95% confidence interval (CI). The 95% CI for the parental Cas12a-expressing strain (PARCas12a) is indicated as a grey dotted line for reference. **(F)** Parasite cell lines were treated with MTF for 48 h and cell densities measured. **(G, H, J)** Parasite cell lines were treated with drug for 48 h and then allowed to recover in drug-free medium for 48-72 h. Cell densities were measured at the end of the treatment phase and after the recovery phase. For all panels, statistical comparisons between mutant strains and the PARCas12a strain were performed using a one-way ANOVA followed by Dunnett’s post-hoc test (significance threshold: p = 0.05; significant differences marked with an asterisk). EC_50_ values for individual addback strains were compared to their corresponding knockout strains using a two-tailed t-test (p = 0.05; significant differences marked with a dagger †).

Instead, resistance correlated more directly with flagellar integrity. Deletion of Arl6/BBS3 (LmxM.16.1380), a BBSome-associated non-core component localized to the flagellum (*86, 87*), also increased Sb^III^ tolerance (fig. 3A) showing modest enrichment in the screen (fig. S6A). Additional resistance-associated genes included cell-body and flagellar-localized proteins UBC2 (LmxM.04.0680), MRK1 (LmxM.31.0120), and PP2C (LmxM.25.0750) (fig. S8A), as well as basal flagellar proteins LmxM.19.0450 (*88*), NPHP4 (LmxM.25.2000) (*89*), and FOPNL (LmxM.32.1895) (*90*) (fig. 2A, S6A). Finally, we identified the glucose transporters LMGT1-3 (LmxM.36.6280-6300), of which LMGT1 (LmxM.36.6300) localizes to the flagellum (*91*), as potential Sb^III^ resistance markers. Enrichment was strongest when sgRNA sets targeted two or all three isoforms, but remained detectable, albeit weaker, for individual isoforms (fig. 2A, S6A). Since FOPNL (LmxM.32.1895) (*90*), NPHP4 (LmxM.25.2000) (*89*), BBS9 (LmxM.31.1240) (*77, 80*) and IFTs (*92*) are known regulators of flagellar/ciliary trafficking, our findings support a model in which Sb^III^ sensitivity depends on an intact flagellum in *L. mexicana*, consistent with mechanisms proposed in macrocyclic lactone action in *C. elegans* in which correct cilia construction is required for drug sensitivity (*78, 79*). This also aligns with earlier reports highlighting diverse flagellar functions in *Leishmania* (*93, 94*) (supplementary discussion 1).

In addition to flagellum-associated genes, a number of other markers were identified. These include ZF-CCCH (LmxM.15.0160), a known Sb^III^ marker (*95, 96*), and DUB22 (LmxM.34.1390), a putative OTU-like cysteine protease with a Zn-finger domain (fig. 2A, S6A, 3A, S4). Of great interest, AQP1 (LmxM.30.0020), a major determinant of Sb^III^ resistance in *L. major* (*14, 18*) and other *Leishmania* species (*17, 19, 20, 23*) did not appear in our screen (fig. 2A, S6A). Furthermore, individual deletion of *L. mexicana* AQP1 did not yield resistance whilst *L. major* AQP1 mutants generated in parallel (fig. 3B, S4) were highly resistant. Species-specific differences in Sb^III^ susceptibility have been reported (*16*) and here we demonstrate a previously unrecognized lack of resistance in AQP1-deficient *L. mexicana*. This is likely due to localisation-differences between species as AQP1 resides in the flagellar membrane of *L. major* (*97*) (and presumably other susceptible species) but is resident in the glycosomal membrane of *L. amazonensis* (*23*), a close relative of *L. mexicana*, hence unable to contribute to uptake from the extracellular environment. This further raises the possibility that *L. mexicana* relies more strongly on broader flagellar-trafficking pathways or alternative membrane transport processes for Sb^III^ uptake, whereas in *L. major*, AQP1 being the dominant uptake route, masks more subtle contributions from other flagellar-associated processes.

**Figure 4.**
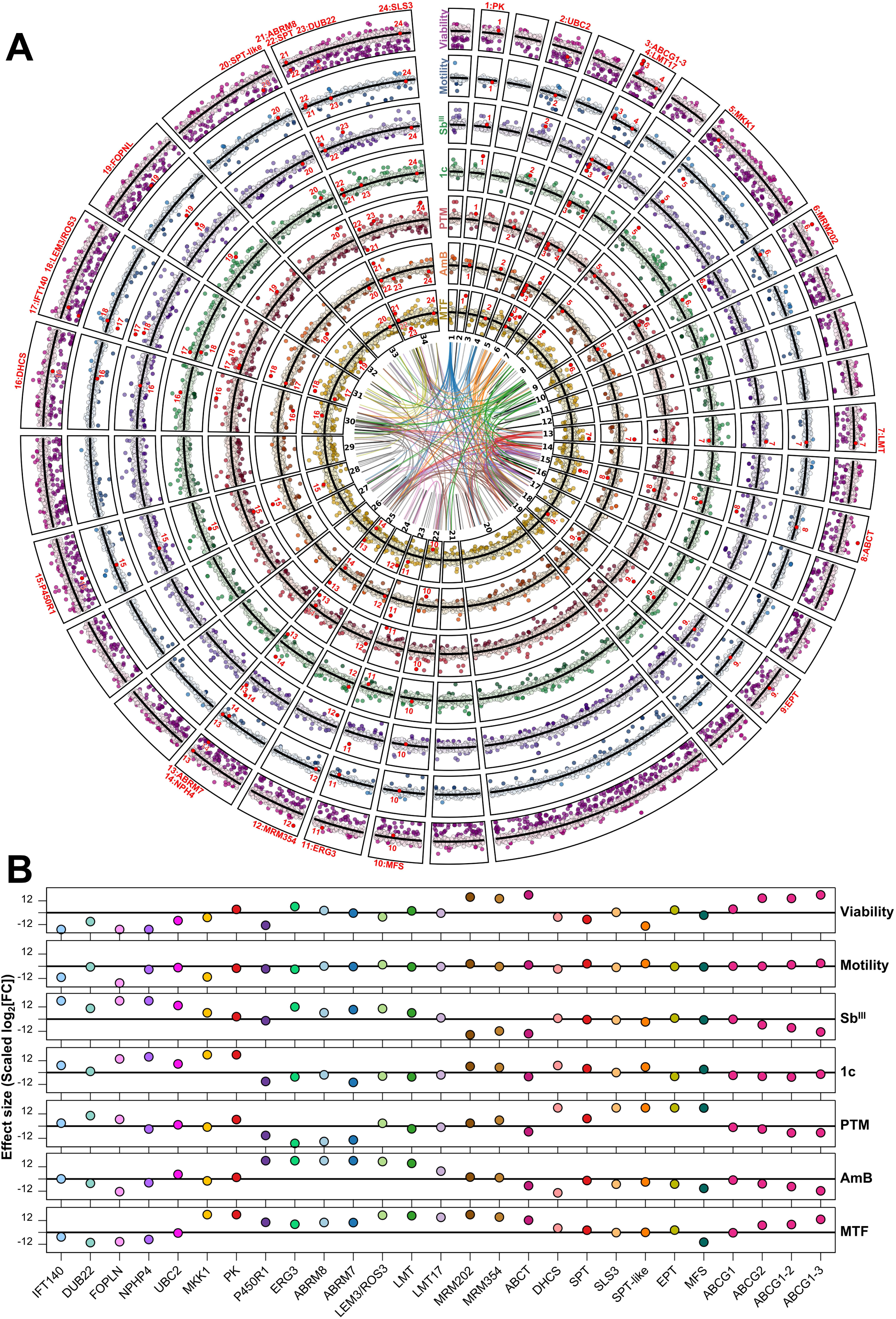
Drug response markers reveal cross-resistance and collateral sensitivities. **(A)** Circos plot displays genome-wide enrichment and depletion of all 7,718 unique or multi-gene sgRNA target sets, mapped to their chromosomal positions. Values represent scaled log_2_ fold changes for the median of each sgRNA set, averaged across three biological replicates. The inner ring represents the 34 chromosomes of *L. mexicana*. Multi-copy sgRNA targets are linked by coloured lines spanning the inner chromosome ring. The seven outer rings show results from (1–5) drug response screens (average log_2_ fold change of 192 h vs. recovery 1 and 264 h vs. recovery 2, referred to as the drug score; supplementary file 7), (6) the motility screen (144 h vs. upper transwell fraction), and (7) the viability screen (0 h vs. 360 h). The black line denotes the baseline (no enrichment or depletion). Data points (sgRNA sets) near the baseline are shown in lighter shades; enriched (above line) or depleted (below line) targets appear as darker-coloured dots (log_2_ fold change ≤1 and ≥1). Genes exhibiting cross-resistance and/or collateral sensitivity are annotated with red numbers and their corresponding gene name is indicated. **(B)** Values as in **(A)** for genes showing cross-resistance and collateral sensitivity.

Interestingly, IFTs, UBC2 (LmxM.04.0680), NPHP4 (LmxM.25.2000), and FOPNL (LmxM.32.1895) were also enriched following 1c treatment (fig. 2B, S6B), further linking altered drug sensitivity to flagellar defects. Individual deletions (fig. 3C & S4) and the screen (fig. 2B, S6B) also showed that mutations in MKK1 (LmxM.08_29.2320) and MPK3 (LmxM.10.0490), either alone or in combination, conferred 1c resistance, consistent with their proposed role in flagellar length maintenance (fig. S8B-C) within a shared signalling cascade (*98, 99*). Additionally, 1c resistance was observed upon targeting another protein kinase (PK, LmxM.02.0570) and an uncharacterized protein, LmxM.10.0940, here named CRM143 (1c resistance marker of 143 kDa) (fig. 2B, S6B, 3C, S4). Conversely, LmxM.20.0690, designated CSM142 (1c hypersensitivity marker of 142 kDa), increased 1c sensitivity in both individual tests and the screen (fig. 2B, S6B, 3C, S4).

Together, these findings reveal both drug-specific and shared resistance mechanisms for Sb^III^ and 1c, highlighting a role for flagellar-associated processes in promastigote sensitivity to particular drugs. The enrichment of multiple independent flagellar assembly components suggests that the flagellum contributes to Sb^III^ and 1c susceptibility, potentially through effects on drug uptake, trafficking, or stress adaptation (supplementary discussion 1). Although the relevance of these mechanisms to clinical resistance remains uncertain given the reduced external flagellum in amastigotes, these findings demonstrate the power of our unbiased genetic screening resource to uncover biological subsystems that become selectively important under anti-leishmanial drug pressure.

### STEROL METABOLISM AND TRANSPORTER FUNCTION SHAPE AMB AND MTF RESPONSE

Overlapping sensitivity was not exclusive to Sb^III^ and 1c, as shared enriched genes were also observed between the 1c and MTF screens. Targeting MKK1 (LmxM.08_29.2320) and PK (LmxM.02.0570) increased resistance to both 1c and MTF (fig. 2B&D, S6B&D, 3C&E, S4). While enrichment of PK and MKK1 decreased with increasing MTF concentrations, enrichment remained significant at lower doses. But in contrast to Sb^III^ and 1c screens, IFT genes were not enriched in any MTF condition (fig. S9A; supplementary discussion 2).

A similar pattern of partial overlap alongside drug-specific mechanisms was also observed between AmB and MTF resistance. As previously reported, mutations in sterol biosynthesis genes conferred AmB resistance (*29–32, 36–40*). Notably, we observed strong enrichment for loss-of-function mutations in the lathosterol oxidase-like protein (LSO-like) ERG3 (LmxM.23.1300) and Lanosterol 14-alpha demethylase (LS14αD) ERG11 (LmxM.11.1100) (fig. 2C&F, S6C), both playing key roles in ergosterol biosynthesis (*31, 32, 38, 40*). In addition, targeting P450R1 (LmxM.28.1240) conferred strong AmB and mild MTF resistance in both the screen and individual deletions (fig. 2C&D, S6C&D, 3D-E, S4). The AmB resistance is consistent with findings showing that P450R1 (needed for ERG11 activity and possibly other cytochrome P450s) mutants lack ergosterol and accumulate 14-methylfecosterol and related intermediates (*37*). P450R1’s role in MTF resistance, though previously unreported, aligns with suggestions linking MTF resistance to disruptions in lipid and sterol metabolism (*41*).

By contrast, ERG2 (LmxM.08_29.2140) showed only mild enrichment with AmB, and other sterol-related genes like ERG4 (LmxM.32.0680), ERG5 (LmxM.29.3550), ERG7 (LmxM.06.0650) and ERG24 (LmxM.31.2320) were not significantly enriched (fig. 2C&F, S6C). We also identified two small hypothetical proteins, ABRM7 (LmxM.25.2375) and ABRM8 (LmxM.34.0270) (AmB resistance marker of 7 and 8 kDa), that conferred AmB resistance in individual deletions and the screen (fig. 2C, 3D, S4, S6C). Both proteins are conserved across the Trypanosomatidae family, with no homologs outside this lineage. Sterol profiling of their null mutants revealed loss of 5-dehydro-episterol alongside increased C28- and C29-diene-sterol-3 levels (fig. S10A-D, supplementary file 8), suggesting a regulatory role in ERG3 (LmxM.23.1300) activity (fig. 2F), and further demonstrating the ability of the library, in conjunction with leishmanicide-selection, to discover new components of biological systems.

Three transmembrane proteins were also enriched in AmB-resistant fractions (fig. 2C, S6C), all of which also conferred strong MTF resistance (fig. 2D, S6D) (fig. 3D&F, S4). Two correspond to known MTF resistance markers, LEM3/ROS3 (LmxM.31.0510) and LMT (LmxM.13.1530), which form the MTF transporter complex (*24–28*). The third is a previously uncharacterized, small hypothetical protein we designate LMT17 (*Leishmania* MTF transporter of 17 kDa; LmxM.06.0995) (fig. 2C-D, 3D-E, S4), which we show is required for the localization of this complex (fig. S11-S13). While earlier reports placed LMT (LmxM.13.1530) at the plasma membrane (*24, 28*), we observed two (fig. S11A-C) and occasionally more (fig. S11D) distinct intracellular signals across multiple C-terminal tags (mNeonGreen (mNG), eYFP, Myc, HA, mCherry (mCh); fig. S11A-D, S12, S13D), resembling localization patterns of other previously reported resistance markers, including PRP1 (*100*), MRPA (*101*), and MDR1 (*102*). Tagging both LMT (LmxM.13.1530) alleles with mNG and mCh does not impair function as MTF resistance isn’t increased (fig. S13A-D). However, deletion of either LEM3/ROS3 (LmxM.31.0510) or LMT17 (LmxM.06.0995) disrupts LMT localisation which is restored by episomal complementation (fig. S13E-G), identifying a hitherto unrecognised requirement for LMT17 in the MTF transporter complex.

Additionally, we identified and confirmed several genes conferring mild MTF resistance, suggesting contributions from pathways beyond the canonical MTF transporter complex (fig. 2D, S6D, S9B, 3E-F, S4; supplementary discussion 3).

### STEROL AND SPHINGOLIPID BALANCE CONTROLS PTM AND AMB SUSCEPTIBILITY

Membrane composition also influenced PTM resistance, but via a distinct mechanism, involving disruption of multiple sphingolipid-metabolism genes, including DHCS (LmxM.30.1780), SPT (LmxM.34.0320), SPT-like (LmxM.33.3740), SLS3 (LmxM.34.4990), EPT (LmxM.18.0810), and others (fig. 2E, S6E, 2G). Individual deletions (fig. S4) increased doubling times (fig. 1E&H, S14A), and confirmed strong PTM-resistance (fig. 3G), although cell-counting was needed as MTT (Thiazolyl Blue Tetrazolium Bromide) dye conversation was impacted for several PTM-resistant lines (fig. S14B). This impairment likely reflects disrupted mitochondrial and redox function, as previously reported for sphingolipid-metabolism mutants (*41, 103–108*), and aligns with mitochondrial exclusion as a mechanism of PTM-resistance in *Leishmania* (*109, 110*). Beyond sphingolipid metabolism, we identified and then confirmed diverse PTM-resistance markers, including genes associated with membrane organization, intracellular trafficking, organelle homeostasis, and post-translational regulation, as well as the adenylate kinase ADK (LmxM.04.0960) and the hypothetical proteins PRM11 (LmxM.33.3955) and PRM60 (LmxM.27.1140) (PTM resistance markers of 11 and 60 kDa) (fig. 2E, S6E, 3G, S4; supplementary discussion 4).

Lipidomic profiling (fig. S10E, supplementary file 9) of selected mutants revealed that DHCS (LmxM.30.1780) and SPT-like (LmxM.33.3740) null mutants exhibited the anticipated (*41, 103–107*) alterations in sphingo-, phospho-, and glycerol-lipid biosynthesis, whereas the ADK (LmxM.04.0960) mutant showed milder shifts limited to phospho- and glycerol-lipids. Notably, lipid changes were minimal in the PRM11 (LmxM.33.3955) null mutant (fig. S10E, supplementary file 9), suggesting that altered lipid composition contributes to PTM resistance, but does not universally explain it.

Interestingly, mutations in sphingolipid metabolism genes that conferred PTM-resistance, including DHCS (LmxM.30.1780), SPT (LmxM.34.0320), SPT-like (LmxM.33.3740), SLS3 (LmxM.34.4990), and EPT (LmxM.18.0810), also increased AmB-sensitivity (fig. 2C, S6C, 3H, S4), despite largely unchanged sterol profiles (fig. S10E, supplementary file 9). This mirrors earlier findings, where disruption of sphingolipid and ceramide biosynthesis compromised protection against ergosterol-binding toxins, increasing AmB-sensitivity (*111*). Conversely, mutations in sterol biosynthesis genes, like ABRM7 (LmxM.25.2375), ABRM8 (LmxM.34.0270), and ERG3 (LmxM.23.1300), conferred AmB-resistance but increased PTM-susceptibility (fig. 2E, S6E, 3I, S4), consistent with reports of sterol mutants exhibiting heightened PTM-sensitivity (*38*). It is intriguing that perturbation of sterol or sphingolipid pathways enhances resistance to one drug while sensitizing the parasite to another, but is likely coincidental reflecting the distinct mechanisms. Elevated mitochondrial membrane potential in *L. major* sterol mutants (*112*) suggested that sterol loss increases PTM sensitivity by enhancing mitochondrial uptake. This is consistent with the mitochondrial exclusion mechanism of PTM-resistance noted above (*109, 110*). Conversely, sphingolipid loss exposes ergosterol to AmB, enhancing its sensitivity.

### COLLATERAL SENSITIVITIES AND OPPOSING DRUG RESPONSES AMONG PARALOGS

Collateral sensitivity-like phenomena extended beyond sphingolipid and sterol metabolism, as additional resistance markers, including multi-copy genes, conferred resistance to one drug while increasing sensitivity to another (fig. 4). For example, targeting DUB22 (LmxM.34.1390) and MFS (LmxM.22.1050) increased MTF sensitivity while conferring PTM resistance (fig. 2A&D, S6A&D, 3G&I, 4). MRM354 (LmxM.24.0260) and MRM202 (LmxM.09.0390), two uncharacterized genes, conferred mild MTF resistance but Sb^III^ hypersensitivity (fig. 2A&D, S6A&D), along with accelerated growth (fig. 1E&H). While no reduced doubling time was observed for Sb^III^-, 1c-, or AmB-response markers, faster growth occurred upon targeting ABCT (LmxM.15.0890) (fig. 1E&H), which conferred low-level MTF resistance (fig. 2D, 3E-F, 4), but increased sensitivity to Sb^III^ and AmB (fig. 2A&C, 3A&C, 4).

The situation with respect to the five other ABCT transporter paralogues identified in the *L. mexicana* genome is intriguing. Two (LmxM.23.0380, LmxM.36.2890), targeted independently, showed no phenotype. In contrast, a sgRNA set targeting all tandem-arrayed ABCG1-3 isoforms (LmxM.06.0080-0100) yielded a similar pattern to ABCT (LmxM.15.0890) (reduced doubling time (fig. 1E, 4), increased MTF resistance, and heightened Sb^III^ and AmB sensitivity (fig. 4, S15)). ABCG transporters are linked to lipid trafficking (*113–115*), thus their disruption may influence drug sensitivities by impairing lipid distribution and membrane architecture.

In addition to these classical collateral sensitivity phenomena, we identified opposing drug responses among paralogous genes for the same compound. For example, ABCC1 (LmxM.23.0210) belongs to a family of seven paralogs, including two tandem arrays (LmxM.23.0210-0220 and LmxM.30.1270-1290). Targeting ABCC1 alone (fig. 2E) or the first array increased PTM resistance, whereas targeting the second array had no effect. Conversely, the seventh paralog (pGP-like; LmxM.30.1430), showed the opposite phenotype, with disruption causing PTM hypersensitivity (fig. 2E, 3I, S4). These findings are consistent with ABCC1-mediated PTM sequestration within target-compartments (*100, 116*) and pGP-like functioning as an efflux pump (*9, 12, 117–119*).

## DISCUSSION

Here, we present a CBE-based loss-of-function screen in *L. mexicana*, targeting 94% of the protein-coding genome to dissect genetic determinants of drug response and fitness in promastigotes. While Cas13b-based (*120*) and RNAi approaches limited to *Leishmania* (*Viannia*) species (*121*) have recently emerged and enable graded knockdown, our system introduces permanent genetic changes, resulting in robust loss-of-function phenotypes. This approach identified both known and previously unrecognized response biomarkers, including systematic evidence linking Sb^III^ sensitivity to flagellar-defects. Functional characterization revealed roles for previously uncharacterized genes, playing roles even in relatively well characterised processes. These include ABRM7 (LmxM.25.2375) and ABRM8 (LmxM.34.0270) in sterol biosynthesis, and LMT17 (LmxM.06.0995) in regulating intracellular localization of the MTF transporter complex. Our data uncovered collateral sensitivities and opposing drug responses in mutants affecting sphingolipid and sterol metabolism, ABC transporters (ABCT, LmxM.15.0890; ABCG1-3, LmxM.06.0080-0100; ABCC1, LmxM.23.0210-0220; pGP-like, LmxM.30.1430) and other genes with diverse functions (MFS, LmxM.22.1050; DUB22, LmxM.34.1390). Uncharacterized loci (MRM354, LmxM.24.0260; MRM202, LmxM.09.0390) also reveal further insights into such phenomena. Further cross-resistance associated genes included DUB22 with Sb^III^/PTM, flagellar-associated genes (IFTs; FOPNL, LmxM.32.1895; NPHP4, LmxM.25.2000; UBC2, LmxM.04.0680) with Sb^III^/1c; PK (LmxM.02.0570) and MKK1 (LmxM.08_29.2320) with Sb^III^/MTF; and P450R1 (LmxM.28.1240), LMT (LmxM.13.1530), LEM3/ROS3 (LmxM.31.0510), and LMT17 (LmxM.06.0995) with AmB/MTF. These findings in *L. mexicana*, a genetically tractable model, support rational combination therapy design, suggesting regimens such as AmB/PTM, MTF/PTM, or MTF/Sb^III^, while cautioning against AmB/MTF, MTF/1c, Sb^III^/PTM, and Sb^III^/1c due to shared resistance profiles.

This is particularly relevant given the current use of MTF/AmB combinations, where emerging cross-resistance could undermine efficacy (*122*), especially if associated with enhanced parasite fitness (*123, 124*). We observed increased fitness in MTF-resistant mutants, including MRM202 (LmxM.09.0390), ABCT (LmxM.15.0890) and MRM354 (LmxM.24.0260). However, drug concentration strongly influenced response profiles, as observed in the MTF screen, indicating that drug responses are context-dependent and clinically relevant amastigote responses are needed too, in order to optimise decisions on combination partners. Species-specific differences may also limit generalization across *Leishmania*, as highlighted here by divergent AQP1-associated phenotypes.

Despite these limitations, many findings likely extend to amastigote stages. For instance, ABCT mutants were enriched during mouse infections (*125*), and sphingolipid-deficient parasites remain infectious by salvaging host lipid precursors (*126–129*). Similarly, PK (LmxM.02.0570), MFS (LmxM.22.1050), and LMT (LmxM.13.1530) mutants showed no detectable amastigote phenotype in previous studies (*61, 125, 130*). For sterol biosynthesis mutants, including AmB-resistant strains, some remain infectious while others lose infectivity (*30, 32, 34, 38, 40*). Certain resistance genes, including P450R1 (LmxM.28.1240) (*37*) and those involved in flagellar formation (MRK1, LmxM.31.0120; MKK1, LmxM.08_29.2320; MPK3, LmxM.10.0490; BBSome, LmxM.31.1240; IFTs), impair amastigote growth and are therefore less likely to contribute to clinical resistance (*61, 67, 98, 99, 125, 131*). Still, flagellar-defective parasites have been isolated from patients (*132*), so their relevance cannot be excluded, underscoring the need for further investigation in amastigotes and across *Leishmania* species.

We therefore now aim to apply our genome-wide CBE libraries and here presented resource to intracellular and axenic amastigotes across species to identify stage- and species-specific response markers. These efforts will also address limitations such as non-deleterious mutations and incomplete editing by incorporating guide activity reporters to measure base editing efficiency (*133*). While even partial disruption is expected to shift sgRNA representation and is accounted for through normalization to non-targeting controls, these improvements will enhance accuracy and resolution. Further developments will also address the lack of inducible editing, as continued editor expression may affect the timing and extent of gene disruption across the population.

Our study provides a comprehensive dataset that can be explored online, while the cryopreservable library offers a resource for future loss-of-function screens across diverse biological processes, including motility, differentiation, host cell immune regulation, drug responses to anti-leishmanial compounds, and many other functional phenotypes. Together, these resources establish a versatile framework for genome-scale functional genomics in *Leishmania*.

## ONLINE METHODS

### Cell culture and parasite strain

*L. mexicana* (WHO strain MNYC/BZ/62/M379) and *L. major* (MHOM/IL/81/FEBNI) promastigotes were cultured at 28°C in M199 medium (Invitrogen, cat. no. 31100-019) supplemented with 2.2 g/L NaHCO₃, 0.0025% hemin, 0.1 mM adenine hemisulfate, 1.2 µg/mL 6-biopterin, 40 mM HEPES (pH 7.4), and 20% heat-inactivated fetal calf serum (FCS; Invitrogen, cat. no. 10500064). If required, culture media were further supplemented with 40 µg/mL of each appropriate selection antibiotic: Hygromycin B, Puromycin Dihydrochloride, Blasticidin S Hydrochloride, Phleomycin, and G-418 Disulfate (all sourced from InvivoGen). Doubling times were determined as described previously (*58*).

For the genome-wide CRISPR loss-of-function screen and generation of most knockout lines, we used *L. mexicana* constitutively expressing tdTomato from the 18S rRNA SSU locus (*58*) and harbouring the pTB107 plasmid for expression of the CBE, Cas12a, and T7 RNA polymerase (*59*), hereafter referred to as PARCas12a. This cell line was also employed for single-allele tagging of LMT (LmxM.13.1530) and for dependency assays involving deletions of LEM3/ROS3 (LmxM.31.0510) and LMT17 (LmxM.06.0995).

When Cas12a-mediated deletions were unsuccessful (see further details under “Cas9 and Cas12a-mediated gene replacement for generation of knockout and tagging cell lines”), *L. mexicana* harbouring the pTB007 plasmid for expression of Cas9 and T7 RNA polymerase (*134*), referred to as PARCas9, was used instead. This Cas9-expressing line was also utilized for tagging of UBC2 (LmxM.04.0680), MRK1 (LmxM.31.0120), PP2C (LmxM.25.0750), and double-allele tagging of LMT (LmxM.13.1530). *L. major* harbouring the pTB007 plasmid was also used to generate the AQP1 (LmjF.31.0020) knockout.

### CBE sgRNA library design and generation

CBE sgRNA sequences for the genome-wide library were obtained from https://www.leishbaseedit.net (*58*). For each protein-coding gene in the *L. mexicana* MHOM/GT/2001/U1103 genome (TriTrypDB v59) (*135, 136*), all sgRNAs predicted to introduce a STOP codon within the first 50% of the ORF were selected. These were ranked as previously described (*59*), and 1-6 top-ranked sgRNAs per gene were chosen, depending on availability.

After removing duplicates and sgRNAs containing PacI or BbsI restriction sites, 38,422 unique CBE sgRNAs remained, covering 94% of all protein-coding genes. For the remaining 6% of protein-coding genes, no CBE sgRNA could be designed to introduce a STOP codon within the first 50% of the ORF. Overlapping sgRNAs across geneIDs resulted in 7,378 unique gene targets and 340 multi-gene targets. BLAST analysis further classified the multi-gene targets into 190 multi-copy genes (E-value = 0) and 150 isoforms (E-value > 0). Eighteen sgRNA sets targeting unrelated gene pairs were excluded from downstream analysis. This yielded 7,718 total sgRNA sets. Distribution across these sets was: 6.4% with 1 sgRNA, 7.8% with 2, 6.6% with 3, 6.5% with 4, 6.9% with 5, and 65.8% with 6 sgRNAs, averaging ∼5.2 sgRNAs per gene. An additional 1,000 non-targeting sgRNAs (*137*) were included as controls and verified by BLAST to lack matches in currently available *Leishmania* genomes (*136*). The final 39,422-sgRNA library was synthesized by GenScript and cloned into pTB105, optimized for CBE expression in *Leishmania* (*59*).

### Transfection of CBE sgRNA expression library and CRISPR screen

To transfect the CBE sgRNA expression library into *L. mexicana*, we upscaled our previously established Cas12a-based protocol using crRNA-4 to mediate integration into the 18S rRNA SSU locus (*59*). For crRNA synthesis, 40 µl each of 100 µM Cas12a common forward primer (5′ gaaattaatacgactcactataggTAATTTCTACTGTCGTAGAT 3′) and crRNA-4 reverse primer (5′ ttccccgtgttgagtcaaatATCTACGACAGTAGAAATTA 3′) were mixed with 420 µl ddH_2_O and 500 µl FastGene Optima HotStart Ready Mix (Nippon), and amplified in a pre-heated thermocycler: 95°C for 3 min, 35 cycles of 95°C for 15 s, 65°C for 15 s, 72°C for 10 s, and final extension at 72°C for 7 min. For CBE sgRNA library preparation, 480 µg of pTB105-cloned CBE sgRNA library was digested with PacI overnight at 37°C across 8 × 300 µl reactions (60 µg DNA/reaction). Cas12a crRNA and digested library DNA were verified via agarose gel, ethanol-precipitated, and resuspended in ddH_2_O (crRNA: ∼680 µg in 160 µl; linearized library DNA: ∼320 µg in 320 µl). Libraries were divided into triplicates (∼107 µg library DNA/replicate), and 100 ng was separated for 0h NGS analysis. Then, each replicate was supplemented with 226 µg crRNA.

Each triplicate was transfected with 1.5 × 10^9^ PBS-washed promastigotes across 15 electroporation cuvettes (∼7 µg library DNA, ∼15 µg Cas12a crRNA, 1 × 10^8^ cells per cuvette, total of 110 µl) using Lonza’s Basic Parasite Nucleofector Kit and program X-001 on an Amaxa Nucleofector 2b (Lonza). Cuvettes were rinsed 4× with medium and pooled into 750 ml/replicate (2 × 10^6^ cells/ml). After 12 h, selection was initiated using 50 µg/ml Hygromycin B and 50 µg/ml Puromycin Dihydrochloride. The transfection efficiency was determined as previously described (*58*), yielding approximately one transfectant per 70 electroporated cells. This corresponded to a representation rate of ∼500 cells per sgRNA within each independently transfected library replicate.

During the viability screen, cultures were maintained with 50 µg/ml Hygromycin B and 50 µg/ml Puromycin Dihydrochloride for 360 h with regular subculturing (fig. 1B) at ≥2 × 10^7^ cells to retain coverage (≥500 parasites/sgRNA). Given the episomal expression of the CBE, continuous selection was considered the most stable condition to ensure uniform expression across the population and avoid biases from random episome loss in some cells, although prolonged CBE expression may permit residual Cas9 binding at target or edited loci. Importantly, this ensures uniform timing of mutagenesis across all sgRNAs, while controlling for potential biases using randomly distributed non-targeting sgRNAs expressed at the same levels as targeting guides, enabling consistent assessment of sgRNA effects across the library.

At 144 hours post-transfection, 2 × 10^7^ cells per triplicate were placed in the bottom chamber of 4 transwells (24-well insert, 5.0 µm PET clear; cellQart 9325012). After 6 hours of incubation, DNA was extracted from cells that migrated to the upper chamber. In parallel, 2 × 10^7^ cells per triplicate were subcultured into 50 mL complete M199 medium (without addition of Puromycin Dihydrochloride and Hygromycin B) treated with the following concentrations of anti-leishmanial drugs: 5x EC_50_ Sb^III^ (272µM) (230057-10G Sigma), 3.5x EC_50_ AmB (105nM) (A9528-50mg Sigma), 1.0, 1.5 and 2.5x EC_50_ MTF (21.5µM, 32.25µM and 53.75µM) (M5571-50mg Sigma), 5x EC_50_ PTM (3.35µM) (P0547-100mg Sigma) and 10x arylmethylaminosteroid 1c EC_50_ (27µM) (*75*). At 192 hours post-transfection, drug-treated cells were centrifuged, the supernatant removed, and cell pellets resuspended in drug-free complete medium. After 48 hours of recovery, cells were subcultured again as shown in fig. 1B.

For each time point, DNA was extracted from ≥5 × 10^7^ cells using either FastGene Blood & Tissue gDNA Extraction Kit (Nippon FG-70250) or DNeasy Blood & Tissue Kit (Qiagen 69504). Then, 2 µg genomic DNA from each time point and 10 ng DNA from plasmid libraries were amplified using FastGene Optima HotStart Ready Mix (Nippon). Amplification employed standard desalted p5 and p7 primers (Sigma, supplementary file 10), containing inline and i5/i7 barcodes for multiplexing, as well as adapters for Illumina sequencing. Barcodes had a Hamming distance of at least 4 nt. To avoid over-amplification, samples isolated from transfected populations were amplified using 26 PCR cycles, while plasmid samples were amplified using only 16 PCR cycles (ideal number of PCR cycles was assessed by testing 12, 14, 16, 18 and 20 PCR cycles for plasmid samples and 22, 24, 26, 28, 30 and 32 PCR cycles for samples isolated from transfected cells; fig. S1B). Amplicons were pooled in equal ratios, size-selected using SPRIselect beads (Beckman Coulter) and send to Novogene GmbH for partial-lane Illumina sequencing (150 bp paired-end sequencing).

### Analysis of CRISPR screening data

For CRISPR screening analysis, sequencing reads were de-multiplexed in two steps: first using bcl2fastq (Illumina) with i5/i7 indices, and then using cutadapt (*138*) with inline barcodes and a 0.15 error rate. This dual de-multiplexing strategy minimized index hopping. Forward reads were then processed with MAGeCK (*74*) (command: “mageck count -l … -n … --sample-label … --fastq …”) to count and normalize sgRNAs, followed by quality control confirming mapping rates above 80% for all samples, as well as low Gini indices and even read distributions (except for strongly selected and skewed MTF library samples [1.5 and 2.5× EC_50_]) (fig. S1; supplementary file 2).

For PW comparisons (fig. 1C-E, 2, S3, S6-7, S9, S15), we used MAGeCK’s Test function (command: “mageck test -k … -t … -c … -n …”) to calculate sgRNA-level p-values and log_2_ fold changes for each of the 39,422 CBE sgRNAs across all comparisons (supplementary file 3). Gene-level metrics for the 7,718 targets (7,378 unique and 340 multi-copy or isoform gene) were derived using MAGeCK’s default settings, in which p-values are computed using an RRA-based model and log_2_ fold changes are summarized as the median of individual sgRNA-level values (supplementary file 4). Viability samples from 96h, 144h, 192h, 264h, 312h, and 360h post-transfection were compared to the 0h plasmid reference. Motility screens used the upper transwell fraction vs. the 144h library. Drug screen recoveries were compared with their viability screen counterparts (192h vs. recovery 1, 264h vs. recovery 2) (fig. 1B). This pipeline used the default settings of the MAGeCK “count” and “test” commands, which apply median-ratio normalization and false discovery rate (FDR) correction by default (*74*). The --control-sgrna option was not used, as median-ratio normalization does not require predefined control guides and non-targeting sgRNAs showed no systematic bias (except for strongly selected and skewed MTF library samples [1.5 and 2.5× EC_50_]) (fig. S1-3; supplementary file 2). Following this MAGeCK analysis, median sgRNA set- and individual sgRNA-level results from all three independent biological replicates were averaged and standard deviations were determined (supplementary file 3 & 4). Depleted and enriched genes, corresponding to hypersensitivity and resistance markers, respectively, were defined using fold change thresholds of ≤ 0.5 or ≥ 2 (log_2_ of ≤−1 or ≥1), with significance set at p ≤ 0.05 (log_2_ of -4.3219).

For MAGeCK MLE analysis (fig. 1F), we used MAGeCK’s maximum likelihood estimation framework (command: “mageck mle -k … -d … -n … --norm-method median”) to model sgRNA abundance changes across the viability time course. A design matrix including an intercept and a continuous time variable (days post-transfection) was used to estimate time-dependent selection coefficients for each sgRNA. Time z-scores were derived from these coefficients and used to rank sgRNA sets based on their temporal depletion or enrichment. Analyses were performed using the combined dataset of all three biological replicates as well as for each replicate individually. The pooled analysis was used for ranking, while replicate-specific analyses were used to calculate standard deviations (supplementary file 5).

Since the library contained on average 5.2 sgRNAs per unique or multi-gene target, the 1,000 non-targeting controls were grouped randomly into sets of five for both analyses (PW and MLE). While this grouping had little effect on enrichment results (fig. S3B-H), it enabled consistent visualization of targeting vs. non-targeting sgRNA set medians without distortion.

### Rank analysis

For fitness rank plots, log_2_ fold changes from the 360 h versus 0 h plasmid comparison or MLE z-scores were ranked from highest to lowest (PW: fig. 1D; MLE: fig. 1F). Depleted and enriched genes, corresponding to fitness loss and gain, respectively, were defined based on relative enrichment compared to the non-targeting control distribution (outside the 95% population interval of the non-targets; supplementary file 5). For combined drug score rank plots (fig. S6), log_2_ fold changes from drug screen recoveries and their corresponding viability controls (192 h vs. recovery 1, 264 h vs. recovery 2) were averaged across both recoveries (referred to as the drug score; supplementary file 7) and ranked from highest to lowest.

### Replicate concordance analysis

Reproducibility across biological triplicates of the library (fig. S2), in response to drug treatment, transwell selection, or viability screening, was evaluated by assessing replicate concordance of normalized sgRNA counts for depleted and enriched median sgRNA set hits.

For each viability PW comparison (0 h plasmid versus 96 h, 144 h, 192 h, 264 h, 312 h, and 360 h), log_2_ fold changes of median sgRNA sets were ranked, and these sets were classified as depleted or enriched as described in the rank analysis above. For drug-treated and transwell-selected libraries, depleted and enriched median sgRNA sets were defined by thresholds of log_2_ fold change ≤−1 or ≥1 and p ≤ 0.05.

For each identified median sgRNA set, the individual sgRNAs composing the set were retrieved, and their normalized counts were extracted across the three biological replicates for the corresponding condition. Pearson correlation coefficients were then calculated using these normalized counts to assess agreement between replicates, generating a 3 × 3 correlation matrix for each condition.

### Cryopreservation and revival of transfected CRISPR libraries

At 96 hours post-transfection, each biological replicate of the parasite library was cryopreserved in complete culture medium supplemented with 10% DMSO at 1 × 10^8^ cells per vial. Cells were frozen at -80 °C for 24 hours before transfer to liquid nitrogen. For recovery, each parasite library replicate was thawed and seeded at 1 × 10^6^ cells/ml, sub-cultured after 24 hours, and reseeded at 1 × 10^6^ cells/ml to ensure full recovery. Following an additional 24 hours of culture, the libraries resembled the original 144-hour time point of the screen (fig. 1B) and were used for subsequent drug screens as described above. Cryopreservation and revival did not result in loss of library diversity, reduced mapping rates, increased Gini indices, or altered read distributions across replicates (fig. S1F, S2C), indicating that the defrosted library is suitable for a broad range of screening applications without detectable sgRNA bottlenecks. However, the process affected sgRNA enrichment patterns, leading to lower correlation between revived and original libraries compared to within-cohort samples (fig. S2C). Importantly, as revived libraries were re-sequenced and analysed using pairwise comparisons with their corresponding conditions, these shifts did not impact screen outcomes. While revived libraries may show reduced suitability for subsequent promastigote fitness screens due to loss of essential genes during freezing, they remain well suited for comparative and condition-specific screening applications.

### Gene ontology analysis

To examine functional patterns of gene essentiality (fig. 1E, G), GO annotations were retrieved for the *L. mexicana* MHOM/GT/2001/U1103 genome (TriTrypDB v59) (*135, 136*), yielding 2,242 unique GO terms. Genes without GO annotations were categorized as “Unclassified.”

As described in the rank analysis, log_2_ fold changes of median sgRNA sets from the 360 h versus 0 h plasmid comparison were ranked from highest to lowest, and classified as depleted or enriched based on relative enrichment compared to the non-targeting control distribution (outside the 95% population interval; fig. 1D-E; supplementary file 5). Sets not classified as depleted or enriched were considered unchanged.

For GO term ranking (fig. 1G), only terms containing at least 10 genes were included, and the percentage of depleted genes within each term was calculated. For sgRNA sets targeting multiple gene IDs, if all associated sgRNA sets fell within a single fitness category (e.g., depleted), each gene ID was counted once for that category. If sgRNA sets spanned multiple categories (e.g., gene ID 1 depleted and gene ID 2 unchanged), each gene ID was counted once in each respective category. GO terms were then ranked based on the proportion of depleted genes to identify functional classes associated with promastigote fitness.

### Cas9 and Cas12a-mediated gene replacement for generation of knockout and tagging cell lines

To generate knockout or tagging lines, we used the LeishGEdit approach and previously established primer design strategies targeting the 5′ and 3′ UTRs of each ORF, employing either Cas9 sgRNAs (*134, 139*) or Cas12a crRNAs (*59*). Gene candidates selected for experimental validation were chosen on a case-by-case basis, considering factors such as ranked enrichment (p-value, log_2_ fold change, raw guide counts, FDR), reproducibility across replicates, biological plausibility, considering existing literature and relevance to known pathways. Importantly, our dataset enables the investigation of a substantially larger number of candidate genes than those selected here for individual knockouts.

Donor DNA was generated by PCR-amplifying pTBlast and pTPuro cassettes for knockouts or pPLOT cassettes for tagging, using primers with 25-30 nt homology flanks adjacent to the Cas9 or Cas12a target sites. PCR products were generated and transfected into *L. mexicana* as previously described (*58, 59, 134, 139*). Cas12a was primarily used, however, Cas9 was employed when Cas12a failed to produce viable mutants. Primer sequences are provided in supplementary file 10, along with annotations indicating whether Cas12a or Cas9 was used to achieve successful gene deletion.

Successful knockouts and tag insertions were verified by genomic DNA extraction (*58, 140*) followed by PCR amplification of the targeted loci using ORF-spanning primers (primer design as previously described (*65*)), with PF16 (LmxM.20.1400) serving as a positive control. LMT-tagged lines were further validated by Sanger sequencing (fig. S9). Most gene deletions were obtained in non-clonal populations using two selectable markers but clonal lines were required to generate complete null mutants for certain targets, including SLS3 (LmxM.34.4990), MFS (LmxM.22.1050), COPE (LmxM.31.1730) and UBC-like (LmxM.21.0440). Due to a limited number of selectable markers, cloning was also necessary for LMT tagging and dependency experiments.

Addback constructs were generated by PCR amplification, using two ORF-spanning primers containing SpeI and AflII restriction sites, with 2xMyc tags incorporated either in the forward primer for N-terminal tagging or the reverse primer for C-terminal tagging. AQP1 addbacks were generated similarly but without Myc tags. The resulting products were ligated into the pTadd plasmid (*65*) and transfected as episomes, following established protocols (*134*). As episomal expression is known to result in incomplete complementation, addbacks did not always fully restore phenotypes. Cell lines selected for addback generation were chosen randomly, and as all tested lines showed partial or full restoration, addbacks were not generated for every mutant.

### Assessing anti-leishmanial drug response in parental and generated mutant cell lines

EC_50_ values for *L. mexicana* parental and mutant lines (supplementary file 11) were determined using a 96-well plate MTT assay previously described (*58*), with each strain tested in triplicate across various drug concentrations and being incubated with the respective anti-leishmanial for 48 h. Absorbance values were background-corrected using media controls, normalized to no-drug wells, and fitted to a logistic regression model to calculate the EC_50_. Statistical significance was assessed via one-way ANOVA followed by Dunnett’s post-hoc test.

Some mutants exhibited reduced MTT conversion compared to parental cells after incubating 1 x 10^4^, 1 x 10^5^, 1.5 x 10^6^ and 3 x 10^6^ cells with MTT for 4 hours in drug-free medium (fig. S11B). For these lines, log-phase promastigotes were diluted to 4 x 10^5^ cells/ml in 2 ml and treated with 3.5x EC_50_ AmB (105nM) (A9528-50mg Sigma), 2.5x EC_50_ MTF (53.75µM) (M5571-50mg Sigma) or 5x EC_50_ PTM (3.35µM) (P0547-100mg). Cultures were incubated for 48 hours at 28°C with 5% CO_2_ and cell counts were taken at the end of treatment (treatment reads in fig. 3D, 3F, 3H). Cells were then centrifuged, the supernatant discarded, and pellets resuspended in 2 ml of drug-free medium. Parasites were incubated under the same conditions for 48-72 hours (based on recovery rate), and final cell densities were determined (recovery reads in fig. 3D, 3F, 3H).

### Imaging of immunofluorescence-labelled and live-cell tagged lines

Live-cell imaging of *Leishmania* parasites was performed as previously described (*141*). For immunofluorescence staining, ∼2 × 10^6^ cells were washed and resuspended in PBS. Paraformaldehyde was then added to a final concentration of 1.5%, and cells were fixed for 10 min on ice followed by an additional 30-min incubation at room temperature. Fixed cells were washed in PBS, attached by centrifugation onto poly-L-lysine-coated coverslips, and permeabilized for 5 min with 0.25% Triton X-100 in PBS. Coverslips were washed twice in PBS and blocked in 3% BSA (in PBS) for 30 min at room temperature. Primary antibody incubation was carried out for 1 h at room temperature using anti-Myc mouse monoclonal IgG antibody 4A6 (1:500; 05-724, Sigma-Aldrich) and anti-HA mouse monoclonal IgG antibody 12CA5 (1:500), both diluted in 1% BSA. After three PBS washes, cells were incubated with Alexa Fluor 488 donkey anti-mouse IgG secondary antibody (1:500; A21202, Invitrogen) for 1 hour at room temperature. Following three additional PBS washes, coverslips were mounted onto glass slides using DAPI-Fluoromount G (SouthernBiotech).

Both live and stained samples were imaged using a Leica Thunder DMi8 Inverted Widefield Fluorescence Microscope with a ×100 NA 1.30 glycerol immersion objective.

### Sterol analysis

Mid-log phase promastigotes were washed once with PBS and resuspended in 25% m/v KOH dissolved in 3:2 v/v ethanol:water, followed by heating at 85°C for 1 hour. After cooling, samples were partitioned by vortexing for 30 s with an equal volume of n-heptane (Sigma). After allowing layers to separate for 20 min, the upper organic layer was retained as the sterol extract. For gas chromatography-mass spectrometry (GC-MS, performed at MVLS Shared Research Facilities, Mass Spectrometry), sterol was extracted from 3 x 10^8^ parasites in 500 μL n-heptane. Trimethylsilane derivatisation was performed by adding 50 μl MSTFA + 1% TMCS (Thermo Scientific) to 100µL of dried extract, vortexing and incubation at 80°C for 15 min. A retention index was added. GC was performed using 1.0 ml/min Helium carrier gas in a Rxi-5SILMS column (30 m length, 0.25 mm inner diameter, 0.25 μm film thickness, Restek) installed in a Trace Ultra gas chromatograph (Thermo Scientific). Derivatised sample (1 μl) was injected into a split/splitless injector using a surged splitless injection (splitless time 30 s, surge pressure 167 kPa). An initial oven temperature of 70°C was increased to 250°C at a ramp rate of 50°C/min; this was reduced to 10°C/min, with a final temperature of 330°C then held for 3.5 min. Eluting peaks were transferred at an auxiliary transfer temperature of 250°C to an ITQ900-GC mass spectrometer (Thermo Scientific), with an emission current of 250 μA. The ion source was held at 230°C, the full scan mass range was 50–700 m/z with an automatic gain control of 50% and maximum ion time of 50 ms. Blank samples were prepared by heating alcoholic KOH and partitioning with n-heptane. Quality control samples were prepared by pooling equal volumes of all samples. Analysis was performed using Xcalibur Quan Processing 4.7 (Thermo Scientific), with identification either by match to a panel of standards or by comparison to the National Institute of Standards and Technology (NIST) library, also Xcalibur. After calculating initial percentage abundance for different identified peaks, those with a relative abundance of < 0.5% were omitted and percentages recalculated to give the final values shown. For UV spectroscopy, sterol extracts from 8 x 10^7^ cells were measured using a Shimadzu UV-2550 UV-Vis spectrophotometer and UVProbe v2.34 software, using n-heptane as a blank. An ergosterol standard spectrum was obtained using 0.05 mg/ml ergosterol (Sigma) in n-heptane. Biological replicates used relate to sterol extracts prepared independently from separate biological samples. A summary of sterol analysis results is provided in supplementary file 8.

### Lipid analysis

Lipids were extracted from cell pellets according to the method described by Folch et al. (*142*) by the addition of chloroform/methanol (2/1, v/v). The mixture was left to stand at 4°C for 1 h. The samples were partitioned by the addition of 0.1 M KCl and the mixture was centrifuged to facilitate phase separation. The lower chloroform phase was collected, evaporated to dryness under nitrogen gas and reconstituted in methanol. Global lipidomic analysis was performed by high resolution liquid chromatography-mass spectrometry (LC-MS) using a Waters Select Series Cyclic IMS mass spectrometer (cIMS) equipped with a heated electrospray ionization (Z-spray) probe and interfaced with a Waters Acquity Premier liquid chromatography system. Samples (5 µl) were injected onto a Thermo Hypersil Gold C18 column (1.9 µm; 2.1 mm x 100 mm) maintained at 50°C. Mobile phase A consisted of water containing 10 mM ammonium formate and 0.1% (v/v) formic acid. Mobile phase B consisted of a 90:10 mixture of isopropanol-acetonitrile containing 10 mM ammonium formate and 0.1% (v/v) formic acid. The initial conditions for analysis were 65%A-35%B, increasing to 65%B over 4 min and then to 100%B over 15 min, held for 2 min prior to re-equilibration to the starting conditions over 6 min. The flow rate was 400 µl/min. Samples were analysed in positive and negative ionisation modes over the mass-to-charge ratio (m/z) range of 50 to 2,000 at a resolution of 30,000. Fragmentation was achieved using the MSe function of the cIMS with a ramped collision energy of 25 – 45 kV. Raw LC-MS data was processed with Progenesis QI v2.4 software (Non-linear Dynamics). Relative fold quantification was performed by the software using all ion normalization, followed by data filtering based on the ANOVA score (< 0.05), fold change (> 2) and ANOVA FDR (< 0.05). This was performed for data acquired in both positive and negative ionization modes. Significant features were then identified using both the Lipid Maps and HMDB databases with a mass error tolerance of 5 ppm. A summary of lipid analysis results is provided in supplementary file 9.

## AVAILABILITY

All data generated or analyzed during this study are included in the manuscript and supplementary files. CRISPR screening data can be further explored under LeishBASEeditDB, which is an open-source database accessible at https://www.leishbaseeditdb.net.

Libraries and plasmids are available on Addgene (https://www.addgene.org/Tom_Beneke/). Cryopreserved *L. mexicana* libraries are available upon request. Please contact.

## ACKNOWLEDGEMENT

We thank Markus Engstler for providing research resources and infrastructure enabling this project. We thank Joachim Geyer and Isabell Berneburg for providing an aliquot of compound 1c. We thank Eva Gluenz and Andreia Albuquerque-Wendt for helpful discussions regarding Sb^III^ resistance in *L. mexicana* and Jessica Valli for discussions on the BBSome in *L. mexicana*. JA and FL were supported by a Flex fund within the LOEWE Center DRUID (Project D3, B3). TB and NHM were supported by the DFG (project 532631727). Additionally, TB was supported by an EMBO Postdoctoral Fellowship (ALTF 727-2021) and Marie Skłodowska-Curie Actions Postdoctoral Fellowship (101064428 – LeishMOM). Lipid and sterol analysis was carried out at the Glasgow University MVLS Shared Research Facility. MPB is funded by the BBSRC (BB/Y007360/1).

## DECLARATION OF GENERATIVE AI AND AI-ASSISTED TECHNOLOGIES

During the preparation of this work, the author(s) used chatgpt, co-pilot and perplexity for additional literature search and to reduce the number of words in the final manuscript. After using this tool or service, the authors reviewed and edited the content as needed and take full responsibility for the content of the publication.

## SUPPLEMENTARY DISCUSSION

### Supplementary discussion 1. Possible roles of flagellar integrity in Sb^III^ and 1c resistance

The strong enrichment of mutations affecting flagellar-associated genes suggests that flagellar integrity contributes directly to Sb^III^ and 1c sensitivity in *L. mexicana*. One possible explanation is that the flagellum or associated membrane trafficking processes, contribute to drug uptake or sensing in promastigotes. This is supported by the enrichment of multiple independent components linked to flagellar assembly and trafficking, including IFTs, BBS9 (LmxM.31.1240), NPHP4 (LmxM.25.2000) (89), FOPNL (LmxM.32.1895) (90), LMGT transporters (LmxM.36.6280-6300), and signalling-associated kinases involved in flagellar maintenance. The broad representation of functionally connected genes within this pathway argues against isolated effects and instead supports selection acting on a larger biological subsystem.

A similar relationship between ciliary defects and drug resistance has previously been described in *C. elegans* (78, 79), where mutations affecting amphid sensory cilia, IFT complexes, dynein motors, BBSome components, and associated trafficking pathways confer resistance to macrocyclic lactones, including ivermectin and moxidectin. In that system, resistance has been proposed to arise through reduced drug uptake via amphidial structures and/or altered downstream signalling responses linked to ciliary dysfunction. Although *Leishmania* lacks the neuronal signalling architecture of metazoans, the enrichment of numerous independent flagellar assembly components in our screens suggests that the flagellum similarly contributes to drug susceptibility, potentially by regulating access of Sb^III^ and 1c to intracellular targets or by controlling membrane-associated stress adaptation pathways.

The lack of strong resistance phenotypes among mutants affecting general vesicle trafficking or flagellar pocket-associated processes further suggests that resistance is linked more specifically to flagellar integrity rather than to global disruption of endo- or exocytosis. Disruption of IFT or BBSome function may therefore alter trafficking or localization of membrane proteins involved in uptake of Sb^III^ and 1c. Alternatively, flagellar defects may indirectly alter membrane organization, surface composition, or stress adaptation pathways that influence susceptibility to these compounds. This could explain why multiple structurally and functionally distinct flagellar proteins converged on similar resistance phenotypes despite lacking obvious direct connections to drug transport. The additional enrichment under 1c selection of signalling-associated kinases, including MKK1 (LmxM.08_29.2320) and MPK3 (LmxM.10.0490), further supports a possible contribution of flagellum-associated signalling or stress response pathways to drug sensitivity.

### Supplementary discussion 2. Dosing effects of PK and MKK1 disruption on MTF resistance

The enrichment of sgRNA sets targeting PK (LmxM.02.0570) and MKK1 (LmxM.08_29.2320) decreased with increasing MTF concentrations, while remaining significant enriched at lower doses (fig. S9A). This pattern suggests that loss of these kinases confers a selective advantage under moderate drug pressure but becomes less beneficial at higher concentrations. One possible explanation is that MTF exerts dose-dependent effects on distinct cellular processes. At lower concentrations, for example, resistance may reflect altered signalling linked to membrane dynamics and stress response pathways, in which PK and MKK1 are likely involved, potentially modulating the parasite’s ability to adapt to drug-induced stress. At higher concentrations, however, survival in the presence of MTF likely requires mutations conferring strong resistance, involving substantial remodelling of membrane integrity or lipid homeostasis, thereby overriding the more limited protective effects of kinase disruption.

Alternatively, the reduced enrichment at higher drug concentrations may reflect a general fitness cost associated with PK and MKK1 loss, which becomes more pronounced under stronger selective pressure. This is consistent with the absence of enrichment of flagellar-associated genes, including IFTs, in any MTF condition, in contrast to Sb^III^ and 1c screens, suggesting a mechanistically distinct response to MTF.

Together, these observations support a model in which MTF resistance is multifactorial and concentration-dependent, with different genetic determinants contributing under distinct selective regimes.

### Supplementary discussion 3. Additional mechanisms contributing to MTF resistance

In addition to the MTF transporter complex, our screen identified several additional genes conferring mild MTF resistance, including MRM202 (LmxM.09.0390), MRM21 (LmxM.31.3830), ABCT (LmxM.15.0890), and MRM354 (LmxM.24.0260) (fig. 2D, S6D, S9B, 3E-F, S4). MRM202 encodes a large low-complexity protein enriched in oligo-monoamino acid stretches, whereas MRM21 shows weak homology to Cornichon-like trafficking proteins, potentially implicating membrane transport or protein localization pathways in MTF susceptibility.

Interestingly, MRM354 conferred only modest resistance in individual validation assays, detectable primarily through cell-counting approaches, despite being part of a gene cohort that remained enriched in the library under strong selection at 2.5× EC_50_ MTF (fig. S9B). This discrepancy may reflect subtle fitness advantages that become detectable only during prolonged competitive growth under drug pressure. More broadly, the identification of multiple genes with comparatively mild phenotypes suggests that MTF resistance may involve cumulative contributions from diverse cellular pathways.

### Supplementary discussion 4. Diverse mechanisms contributing to PTM resistance

In addition to sphingolipid metabolism defects, our screen identified a broad range of genes contributing to PTM resistance, suggesting that susceptibility to this compound is influenced by multiple partially overlapping cellular processes. These included membrane-associated proteins, such as a putative major facilitator superfamily (MFS) transporter (LmxM.22.1050) and the calcium-motive P-type ATPase PMCA (LmxM.34.2080), as well as factors involved in Golgi-to-ER transport and ER membrane retention, including COPE (LmxM.31.1730) and RER1 (LmxM.22.0580) (fig. 2E, S6E, 3G, S4). The enrichment of these pathways is consistent with the known importance of membrane organization, intracellular trafficking, and organelle homeostasis in PTM activity and resistance.

We also identified post-translational regulators, including a ubiquitin-conjugating enzyme-like protein (UBC-like, LmxM.21.0440) and DUB22 (LmxM.34.1390), the latter of which additionally conferred Sb^III^ resistance (fig. 2A&E, S6A&E, 3A&G, S4), suggesting potential overlap between PTM and antimony stress response pathways. Additional markers included the annotated adenylate kinase ADK (LmxM.04.0960) and the hypothetical proteins PRM11 (LmxM.33.3955) and PRM60 (LmxM.27.1140) (PTM resistance markers of 11 and 60 kDa) (fig. 2E, S6E, 3G, S4), further supporting the genetically heterogeneous nature of PTM resistance. The identification of multiple unrelated resistance-associated genes suggests that PTM susceptibility is not governed by a single dominant pathway but instead reflects the integration of membrane composition, mitochondrial function, intracellular trafficking, and stress adaptation processes.

## SUPPLEMENTARY FIGURE LEGENDS

**Figure S1.**
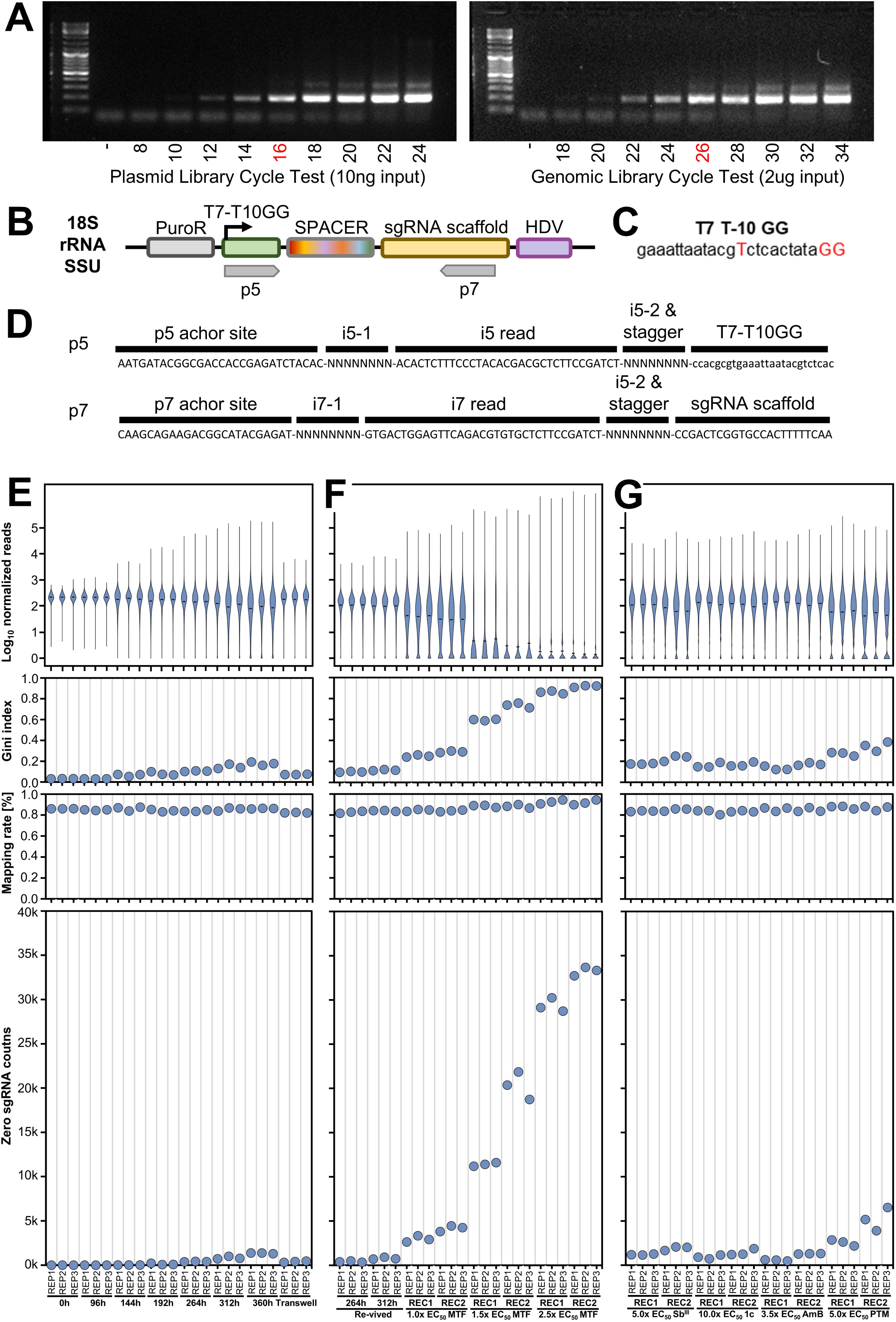
CRISPR screening readout analysis pipeline. **(A)** Determination of the optimal PCR cycle number for Illumina library preparation. A no-template control (–) is included, and the optimal cycle number is indicated in red. **(B)** Schematic of the CBE sgRNA expression construct showing the binding sites for Illumina sequencing primers (p5 and p7). **(C)** Nucleotide sequence of the optimized T7 RNA polymerase promoter used for CBE sgRNA expression (*59*). **(D)** Diagram of p5 and p7 Illumina sequencing primers. **(E-G)** Quality control of individual sgRNAs showing sgRNA-read distributions, Gini indices, mapping rates, and zero sgRNA counts across samples and replicates.

**Figure S2.**
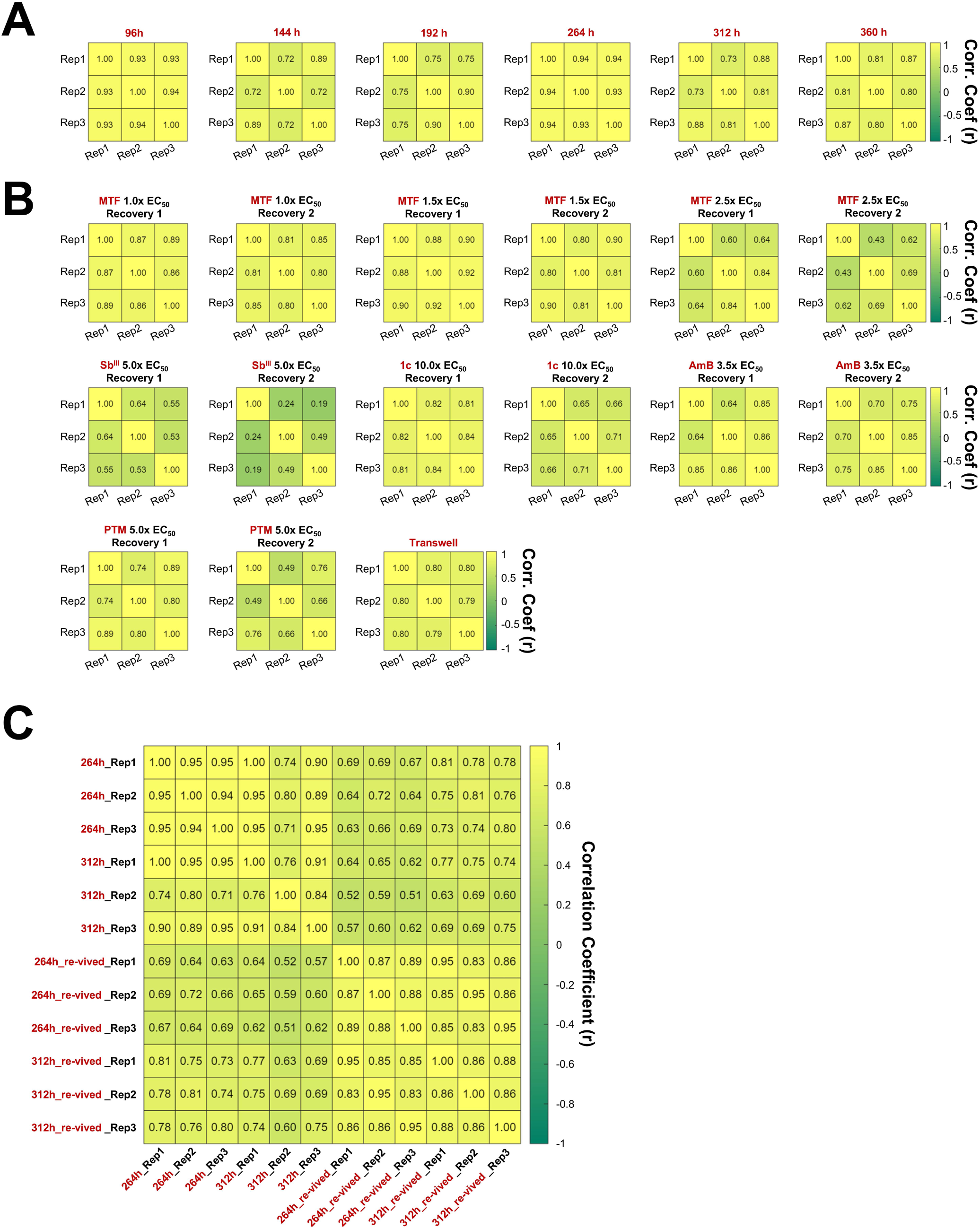
Replicate concordance analysis. Heatmaps show Pearson correlation coefficients (r) among the three biological replicates for depleted and enriched sgRNA hits for **(A)** the viability screen across time points, **(B)** drug treatment and transwell selection across recovery time points, and **(C)** comparison between freshly transfected and revived libraries, considering only shared depleted and enriched sgRNA sets.

**Figure S3.**
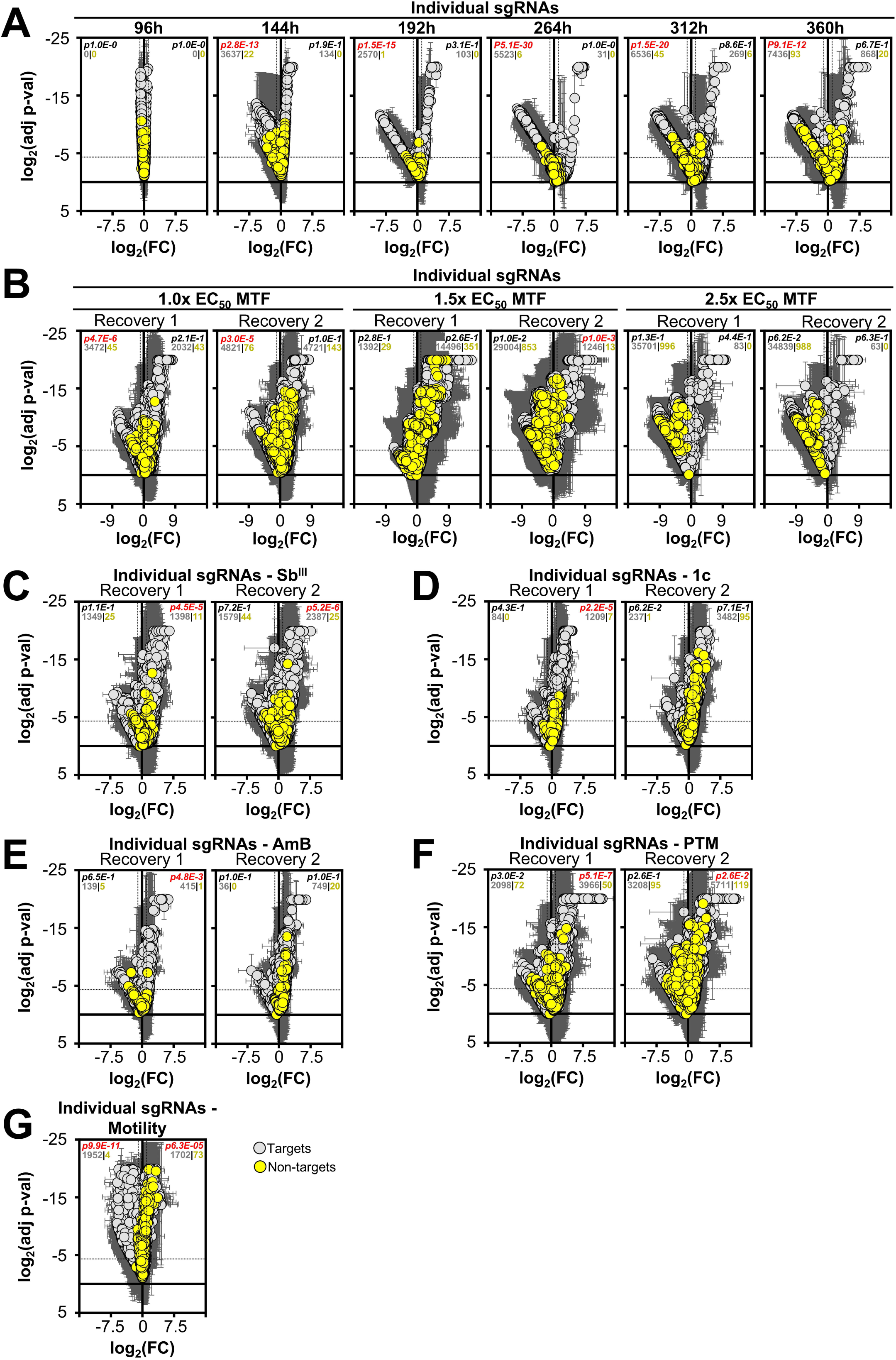
Enrichment of individual sgRNAs. **(A-G)** Distribution of log_2_ fold change (x-axis) and log_2_ p-values (y-axis) for all 38,422 targeting and 1,000 non-targeting sgRNAs. Error bars represent standard deviation across three biological replicates. The number of significantly enriched or depleted targeting and non-targeting sgRNAs (thresholds: log_2_ fold change ≤ -1 or ≥ 1; log_2_ p-value ≤ -4.3219) is indicated in each panel. A chi-square test compared the total of targets (38,422) and number of enriched/depleted targets with the total of non-targets (1,000) and number of enriched/depleted targets. Significant p-values ( ≤ 0.05) are highlighted in red. Comparisons include: **(A)** 0 h plasmid reference vs. post-transfection time points (96 h, 144 h, 192 h, 264 h, 312 h, 360 h), **(B-F)** recovery fractions vs. their matched viability time points (192 h vs. recovery 1; 264 h vs. recovery 2), and **(G)** 144 h post-transfection vs. transwell upper layer (motility screen).

**Figure S4.**
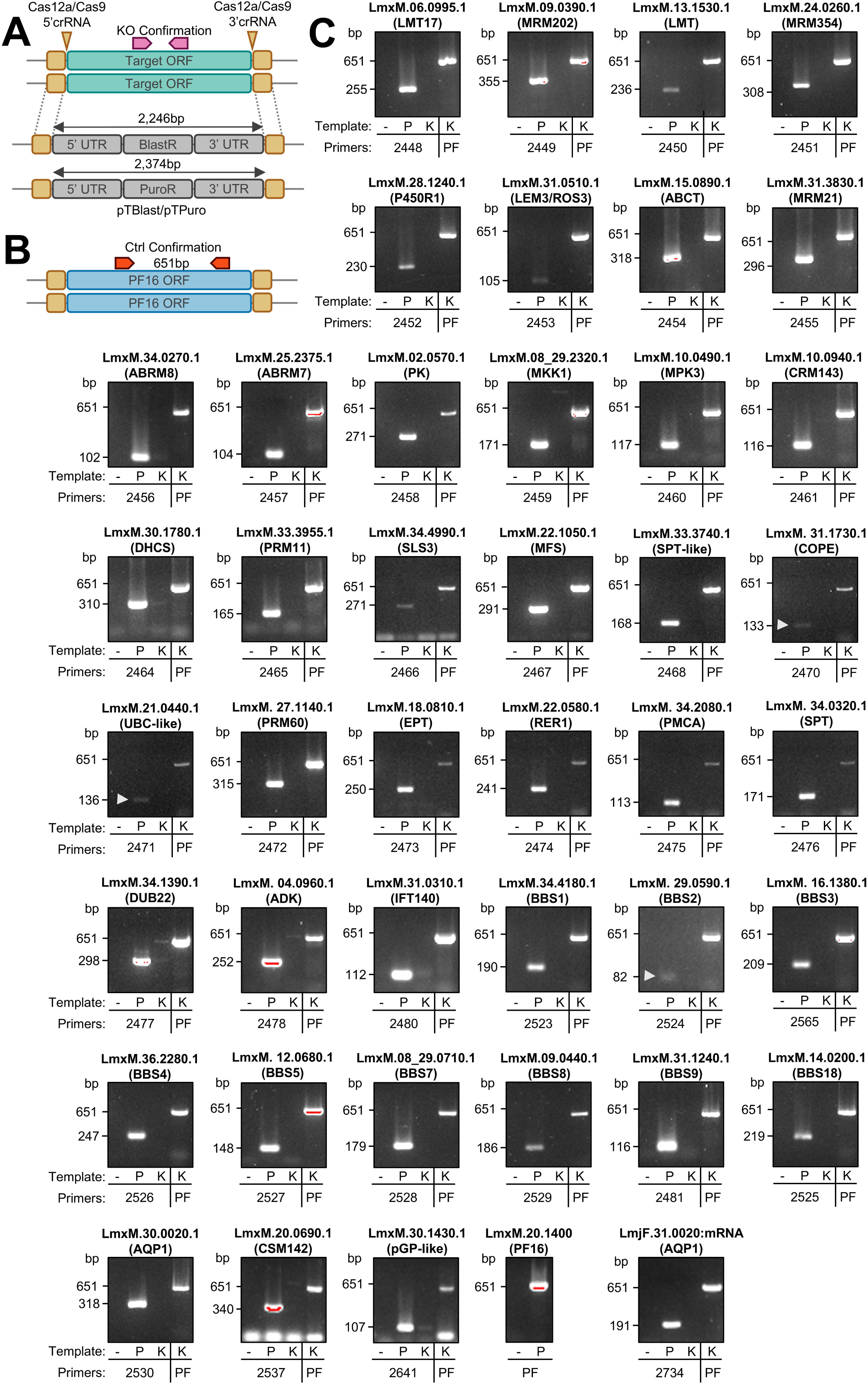
Mutant verification by PCR. **(A)** Schematic of the target locus before and after editing with Cas12a or Cas9, indicating crRNA binding sites, ORF replacement with pTBlast or pTPuro, and locations of primers used to confirm gene knockout. **(B)** Diagram illustrating PCR-based ORF amplification for knockout validation. PF16 primers were used as a positive control to verify quality of genomic DNA. **(C)** Each panel shows PCR amplicons from genomic DNA of the putative knockout line (K), parental strain (P), and a no-template control (-). Genomic DNA from knockout lines was also amplified with PF16 primers (PF) as a positive control. Target gene names and corresponding GeneIDs are indicated. Knockouts were confirmed by absence of the specific ORF amplicon in the mutant. White arrows indicate weak but correctly sized amplicons.

**Figure S5.**
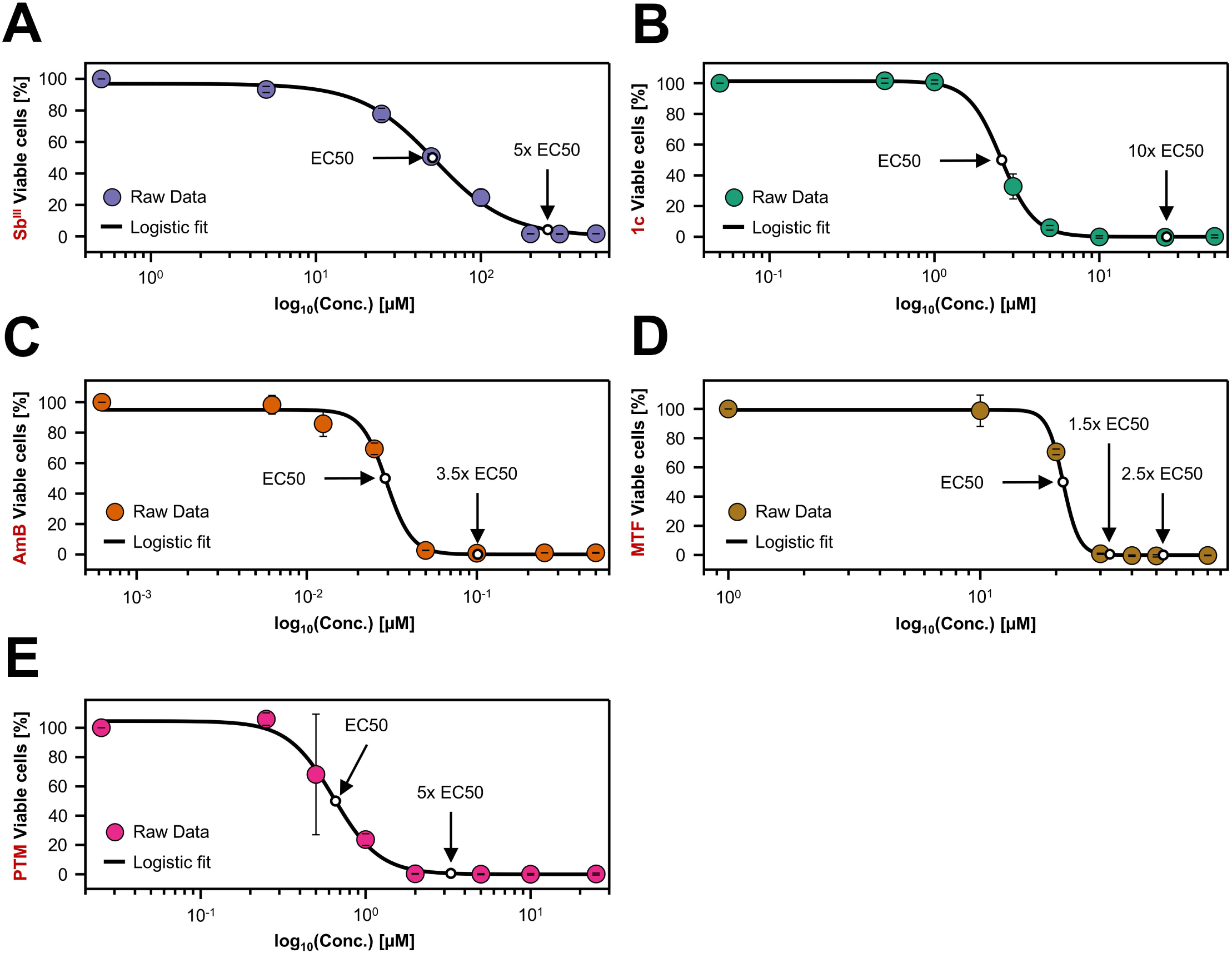
Identification of optimal treatment doses for CRISPR screening in *L. mexicana*. **(A-E)** MTT assay results showing the mean number of viable cells across three independent replicates. Error bars represent standard deviation between replicates. EC_50_ values were calculated by fitting the data to a logistic regression model (black solid lines). The determined EC_50_ values and the drug concentrations selected for the CRISPR screen are indicated on each plot.

**Figure S6.**
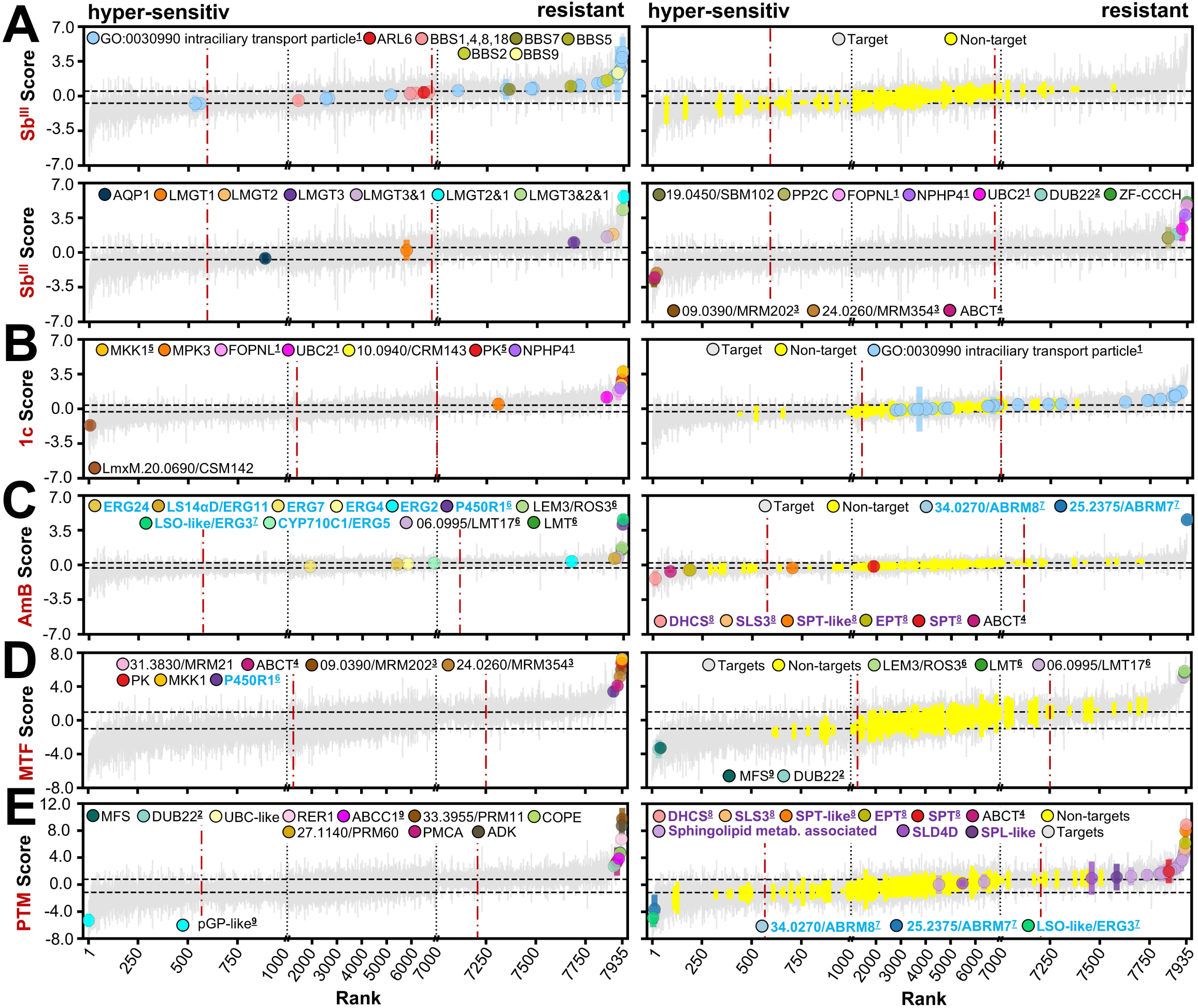
Ranked combined drug score highlight resistance and hypersensitivity markers. Combined scores (log_2_ fold change of recovery 1 and 2 averaged) were ranked for each drug and plotted, showing median sgRNA set values (grey: targeting; yellow: non-targeting), with bars representing standard deviations across triplicates. Horizontal dotted lines indicate significance thresholds (mean log_2_ fold change ≤−1 or ≥1). Vertical dotted black lines denote x-axis zoom regions (first and last 1,000 ranked sgRNA sets), and vertical red dotted lines indicate the 95% reference interval based on non-target distribution. Gene names highlighted in cyan denote sterol metabolism genes, and in purple sphingolipid metabolism genes.

**Figure S7.**
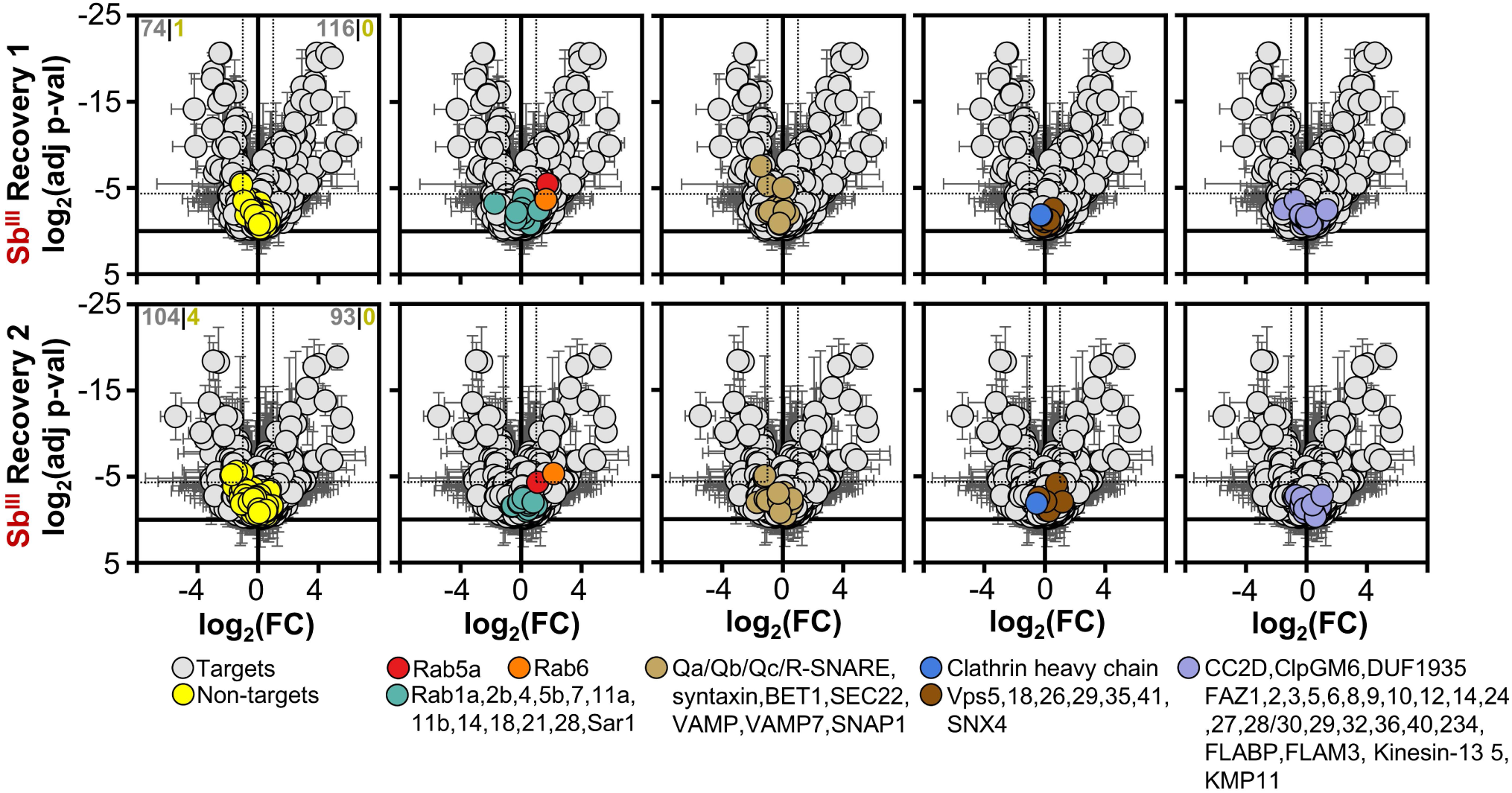
No enrichment of vesicle trafficking or FAZ-associated genes in Sb^III^ drug screen. Median values of sgRNA sets targeting 7,718 unique or multi-gene targets are shown. Error bars represent standard deviations from three independent biological replicates. Comparisons are between Sb^III^ recovery fractions and their corresponding viability screen time points (192 h vs. recovery 1, 264 h vs. recovery 2). Legends below each plot highlight genes involved in vesicle trafficking, grouped from left to right as Rabs, SNAREs, Clathrin, and VPS proteins (*84, 85*), and genes associated with the FAZ (*83*). The number of significantly enriched or depleted sgRNA sets is indicated in the leftmost panel, with non-targeting controls in yellow and targeting sgRNA sets in grey. Significance thresholds used are log_2_ fold change ≤−1 or ≥1, and log_2_ p-value ≤−4.3219.

**Figure S8.**
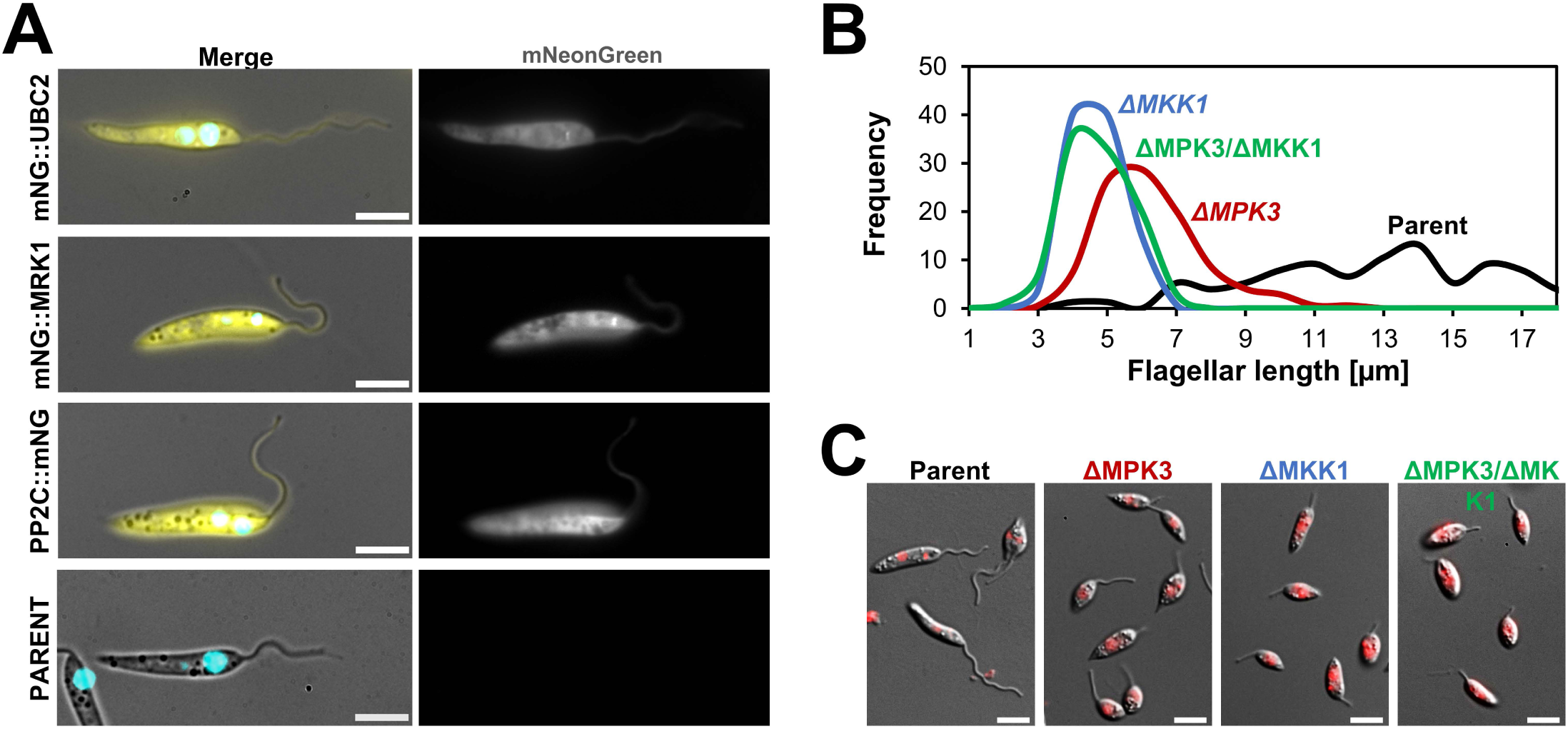
Confirmation of flagellar-associated localizations and phenotypes. **(A)** Representative fluorescence images of N-terminally mNG-tagged UBC2 and MRK1, and C-terminally tagged PP2C. Merged images show DIC overlaid with mNG fluorescence (yellow) and DAPI-stained DNA (cyan). Exposure and imaging settings were normalized and applied equally to both tagged and parental control samples. Scale bar: 5 µm. **(B)** Quantification of flagellar length in ΔMKK1, ΔMPK3, and ΔMKK1/ΔMPK3 mutants, compared to the parental strain. Measurements were based on ≥100 cells per genotype. **(C)** Representative micrographs of parental, ΔMKK1, ΔMPK3, and ΔMKK1/ΔMPK3 cell lines. DAPI-stained DNA is shown in red. Scale bar: 5 µm.

**Figure S9.**
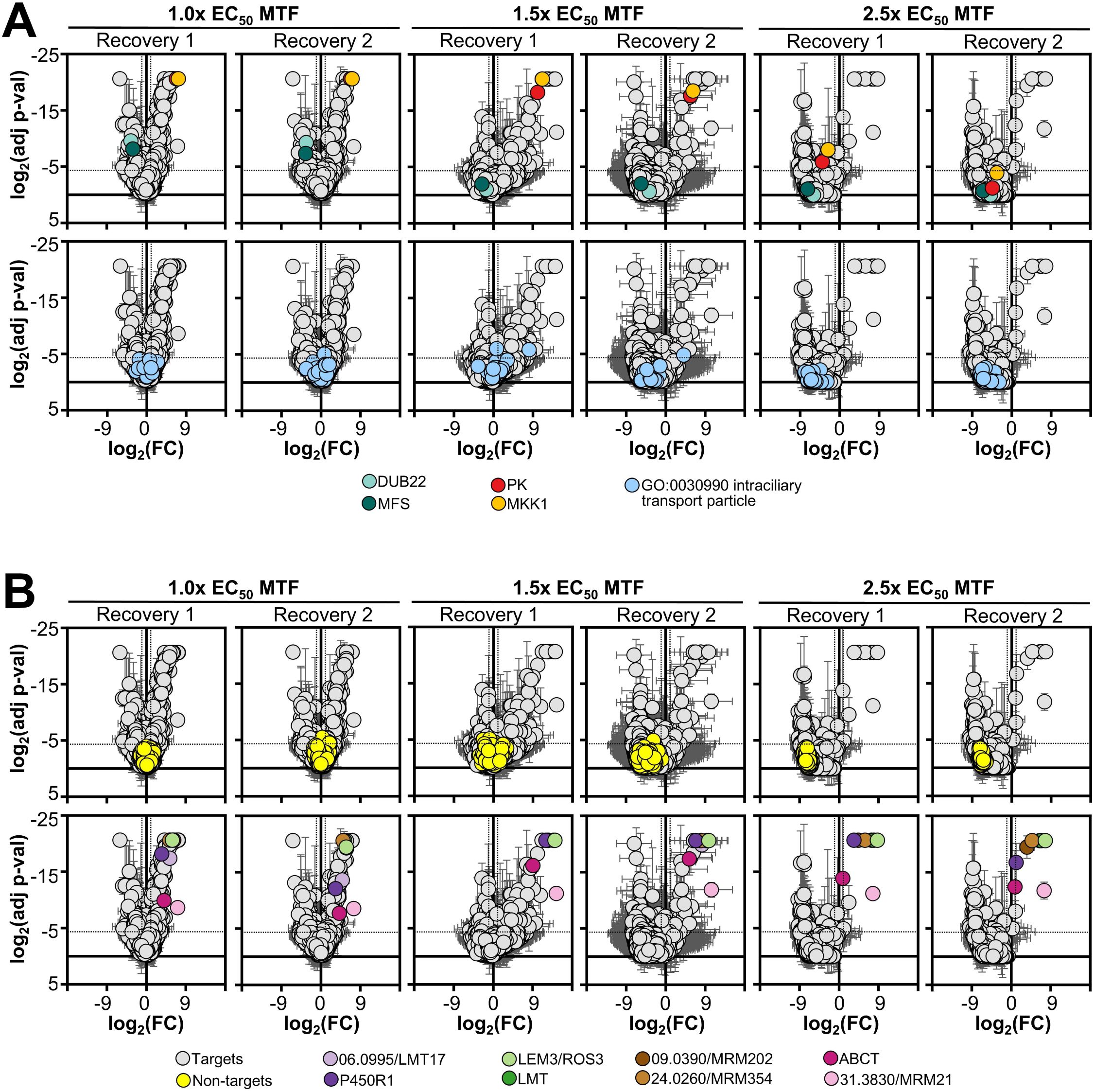
Selected resistance and hypersensitivity markers following MTF screening. **(A-B)** Median sgRNA set values for 7,718 unique or multi-gene targets are shown, with error bars representing standard deviations across three biological replicates. Comparisons are between recovery fractions and corresponding viability time points (192 h vs. recovery 1; 264 h vs. recovery 2), with the MTF screen performed at 1.0×, 1.5×, or 2.5× EC_50_. Selected genes are highlighted in the plot legends.

**Figure S10.**
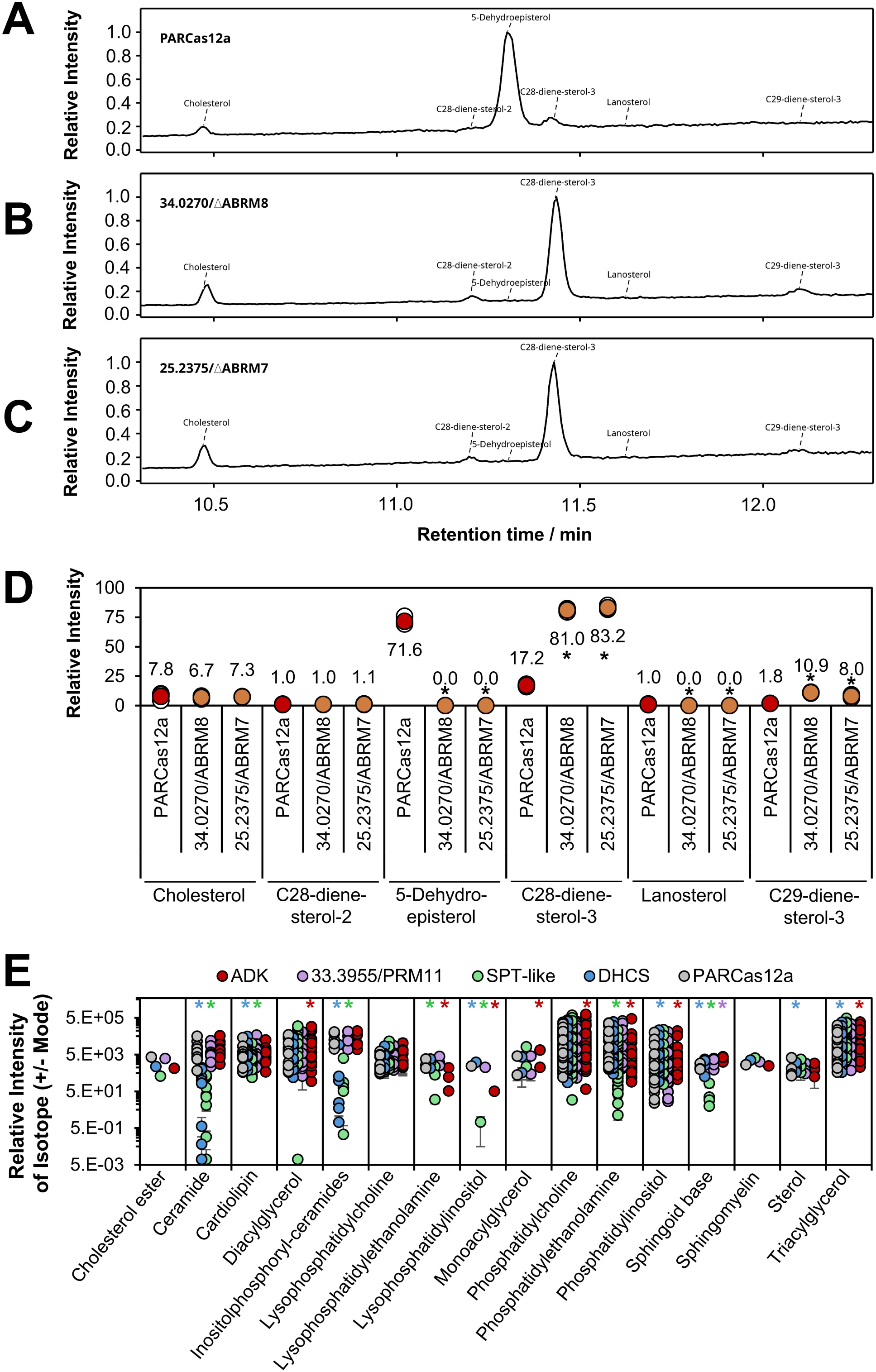
Sterol and lipid-profiling of selected mutant cell lines. **(A-C)** Sterol profiles for ABRM7 (LmxM.25.2375) and ABRM8 (LmxM.34.0270) null mutants and their parental control. Plots show relative intensity vs. retention time (min). Selected sterols are highlighted.**(D)** Quantification of individual sterol isotopes from panels (A-C). Error bars represent standard deviation across three biological replicates. Statistical significance was assessed using a two-tailed t-test (threshold: p = 0.05; significant differences indicated with an asterisk). **(E)** Lipidomic profiling of null mutants for ADK (LmxM.04.0960), PRM11 (LmxM.33.3955), SPT-like (LmxM.33.3740), and DHCS (LmxM.30.1780), along with parental cells. Relative intensities of detected lipid isotopes (in positive and negative ionization mode) are grouped by lipid class and the average of four biological replicates is shown. For each isotope, a Mann-Whitney U test was performed comparing mutant vs. parent. Resulting p-values were combined per lipid class using Stouffer’s method. Significance threshold: p = 0.05; significant differences indicated by an asterisk color-coded according to the corresponding mutant cell line.

**Figure S11.**
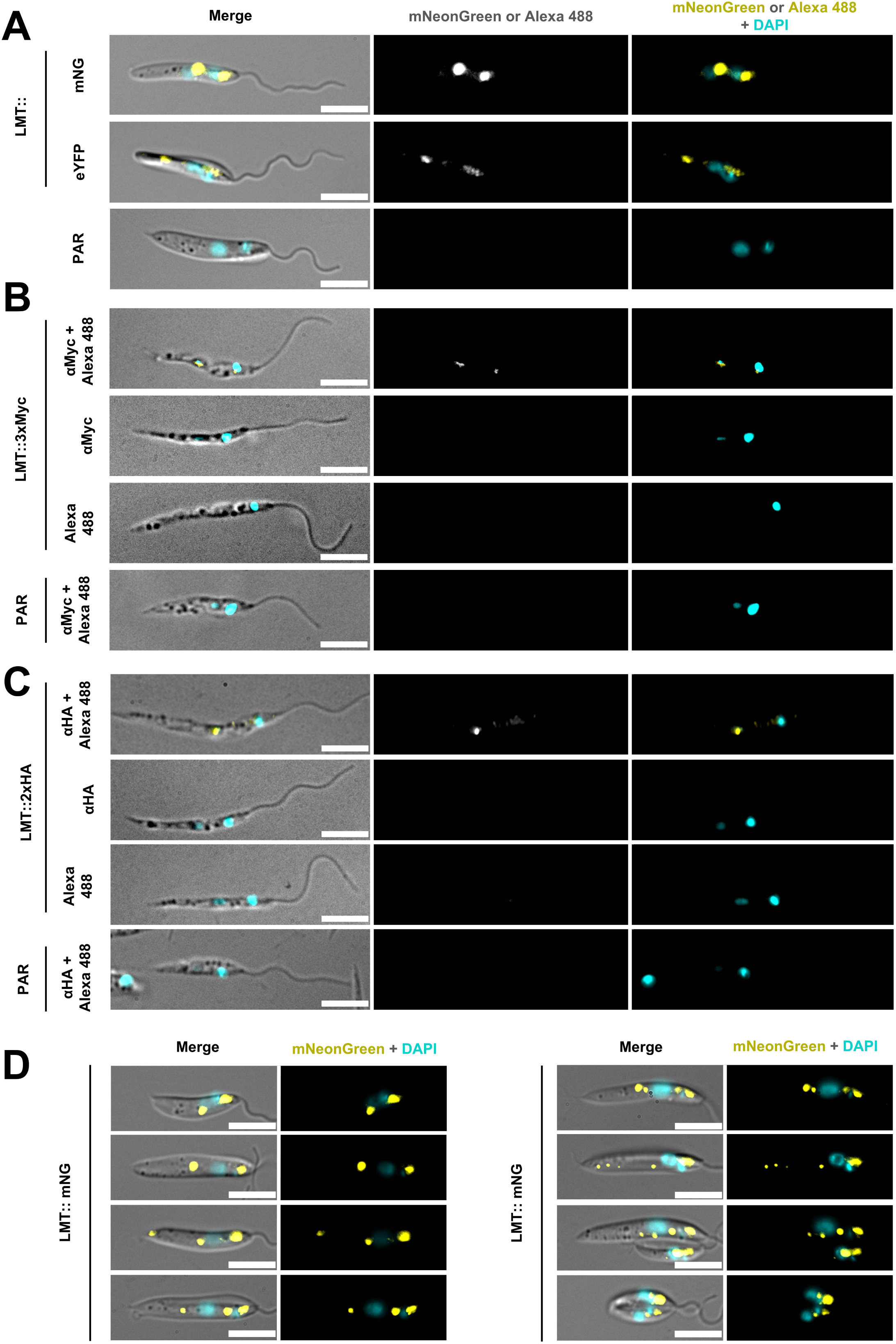
Localization of LMT. **(A)** Representative micrographs of LMT::mNG- and LMT::eYFP-tagged cell lines. Merged images show DIC overlaid with fluorescence signals from mNG or eYFP (yellow) and DAPI-stained DNA (cyan). **(B, C)** Immunofluorescence images of fixed cells expressing LMT tagged with **(B)** Myc or **(C)** HA epitopes. Each panel (top to bottom) shows: cells labelled with both primary and secondary antibodies; cells with primary antibody only; secondary antibody only; and control parental cells stained with both antibodies. Merged images show DIC overlaid with Alexa Fluor 488 signal (yellow) and DAPI-stained DNA (cyan). **(D)** Additional micrographs of LMT::mNG-tagged cell line. Micrograph properties as in **(A)**. Exposure and imaging settings were standardized and applied equally to all tagged and parental control samples. Scale bar: 5 μm.

**Figure S12.**
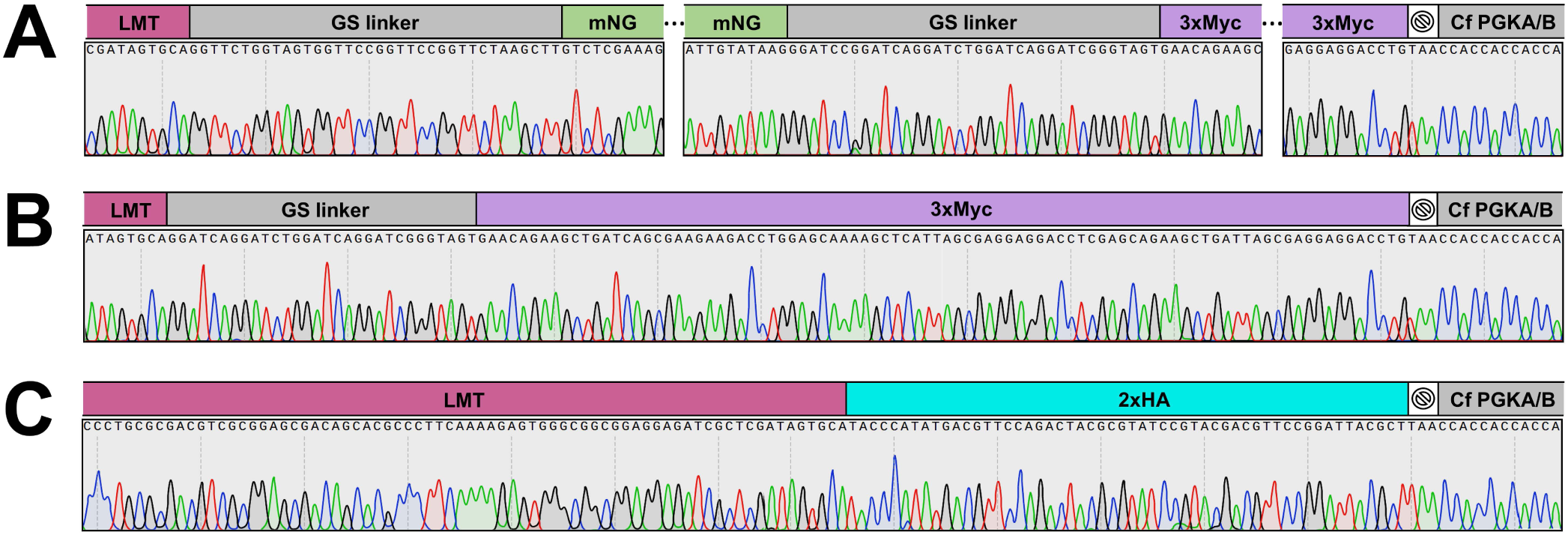
Verification of LMT tagging by Sanger sequencing. The LMT coding sequence (CDS) was C-terminally tagged, and correct integration of the tags was confirmed by Sanger sequencing for: **(A)** mNG, **(B)** 3×Myc, and **(C)** 2×HA.

**Figure S13.**
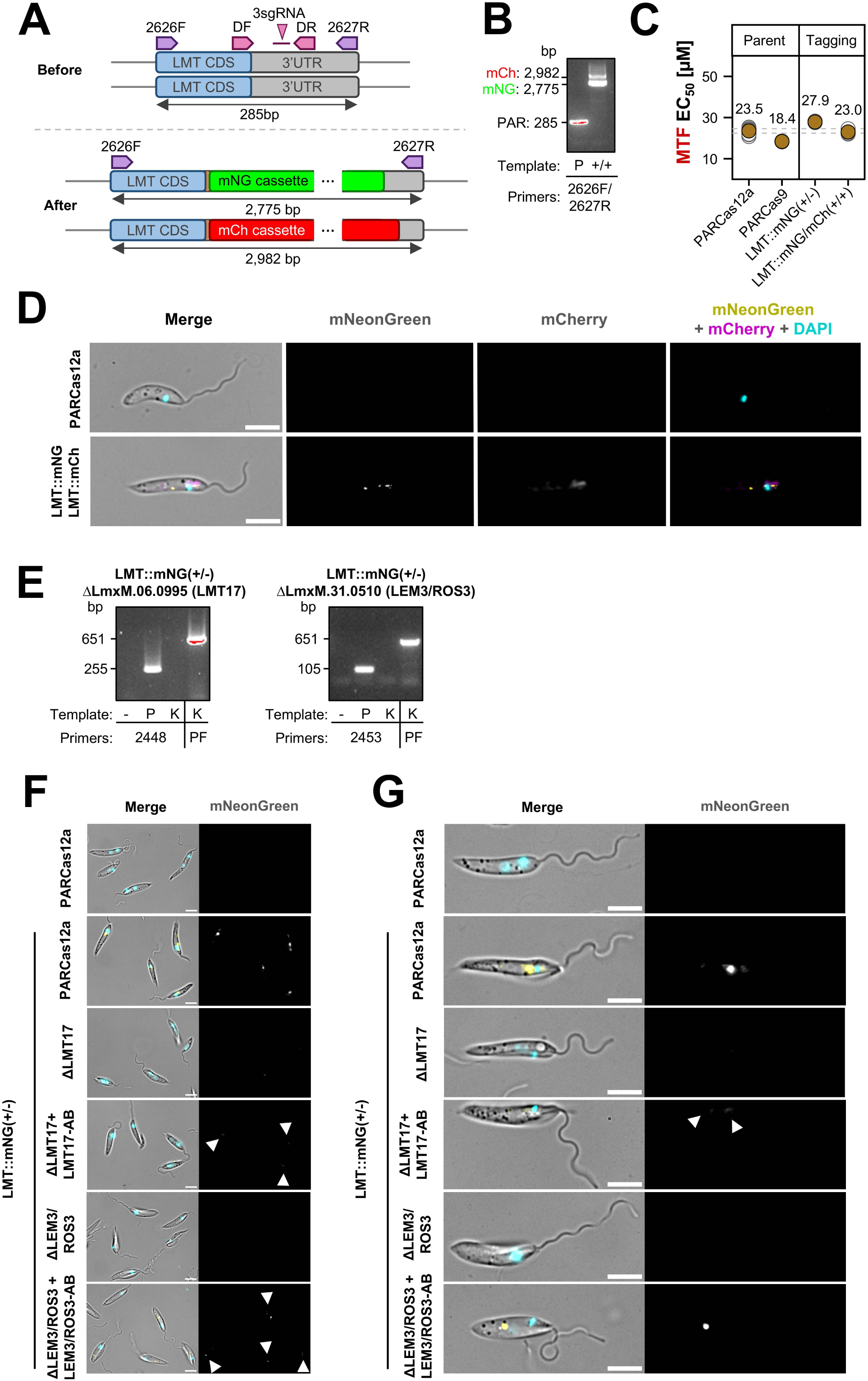
LMT localization depends on LEM3 and LMT17. **(A)** LMT was C-terminally tagged with both mCh and mNG. Tagging of both alleles in clonal cell lines was confirmed by PCR. Diagrams illustrate the LMT locus before and after tagging, with primer positions and expected amplicon sizes indicated. **(B)** PCR products shown correspond to the schematic in **(A)**, confirming the presence of the unmodified locus in the parental cell line (P), and successful integration of both tags with complete modification of the wild-type locus in the double-tagged LMT::mNG/mCh line (+/+). **(C)** EC_50_ values are shown for LMT-tagged cell lines, mNG-tagged either on a single allele using Cas12a (+/-) or mNG/mCh-tagged on both alleles using Cas9 (+/+), alongside their respective parental controls (PARCas12a and PARCas9). Each filled-coloured dot represents the mean EC_50_ from up to nine biological replicates (grey unfilled dots), with error bars indicating the 95% confidence interval (CI). The 95% CI for PARCas12a is marked by a grey dotted line. No significant differences were observed. **(D)** Representative micrographs of the LMT::mNG/mCh double-tagged cell line. Merged images show DIC overlaid with fluorescence signals from mNG (yellow), mCh (pink) and DAPI-stained DNA (cyan). **(E)** LMT::mNG (+/-) cells were subjected to LEM3/ROS3 or LMT17 deletion. PCR analysis shows amplicons from genomic DNA of knockout (K), parental (P), and no-template control (-). PF16 primers (PF) were used as a positive control. Gene names and GeneIDs are indicated. Knockouts were confirmed by loss of ORF-specific PCR bands. **(F-G)** Live-cell imaging of LMT::mNG (+/-) lines following deletion of LMT17 or LEM3/ROS3, and their respective addbacks (LEM3/ROS3-AB and LMT17-AB). **(F)** Multiple cells and **(G)** selected single cells are shown in DIC merged with fluorescence signals from mNG (yellow) and DAPI-stained DNA (cyan). White arrows highlight partial restoration of signal in LEM3/ROS3 and LMT17 addbacks. Exposure settings were standardized across samples. Scale bar: 5 μm.

**Figure S14.**
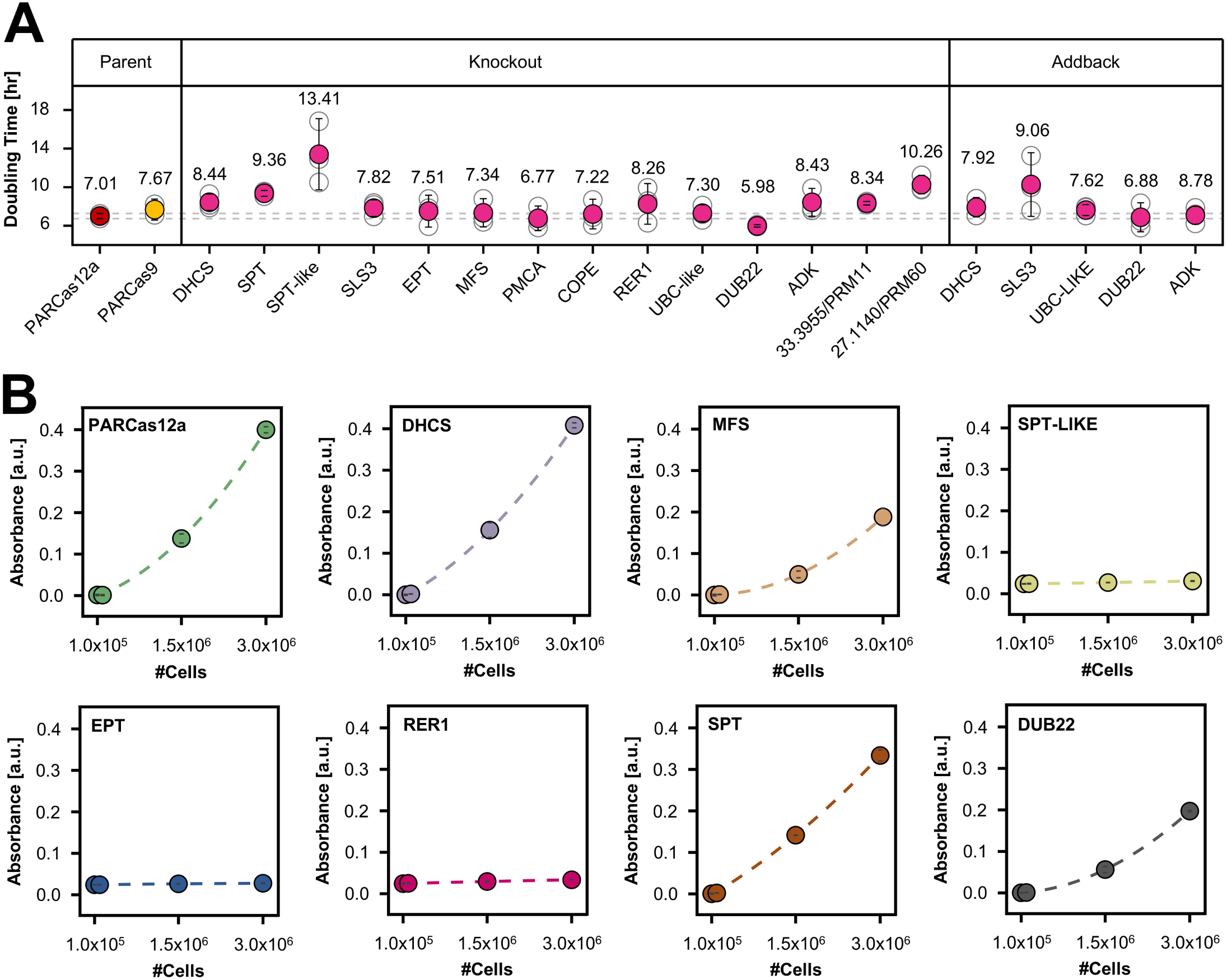
Characterization of knockout cell lines associated with PTM resistance. **(A)** Doubling times of generated null mutants linked to PTM resistance. Error bars indicate standard deviation from three biological replicates. **(B)** PTM-associated mutants and their parental control (PARCas12a) were incubated with MTT for 4 h using 1 x 10^4^, 1 x 10^5^, 1.5 x 10^6^, and 3 x 10^6^ cells per well. Absorbance values were measured and plotted against the number of plated cells.

**Figure S15.**
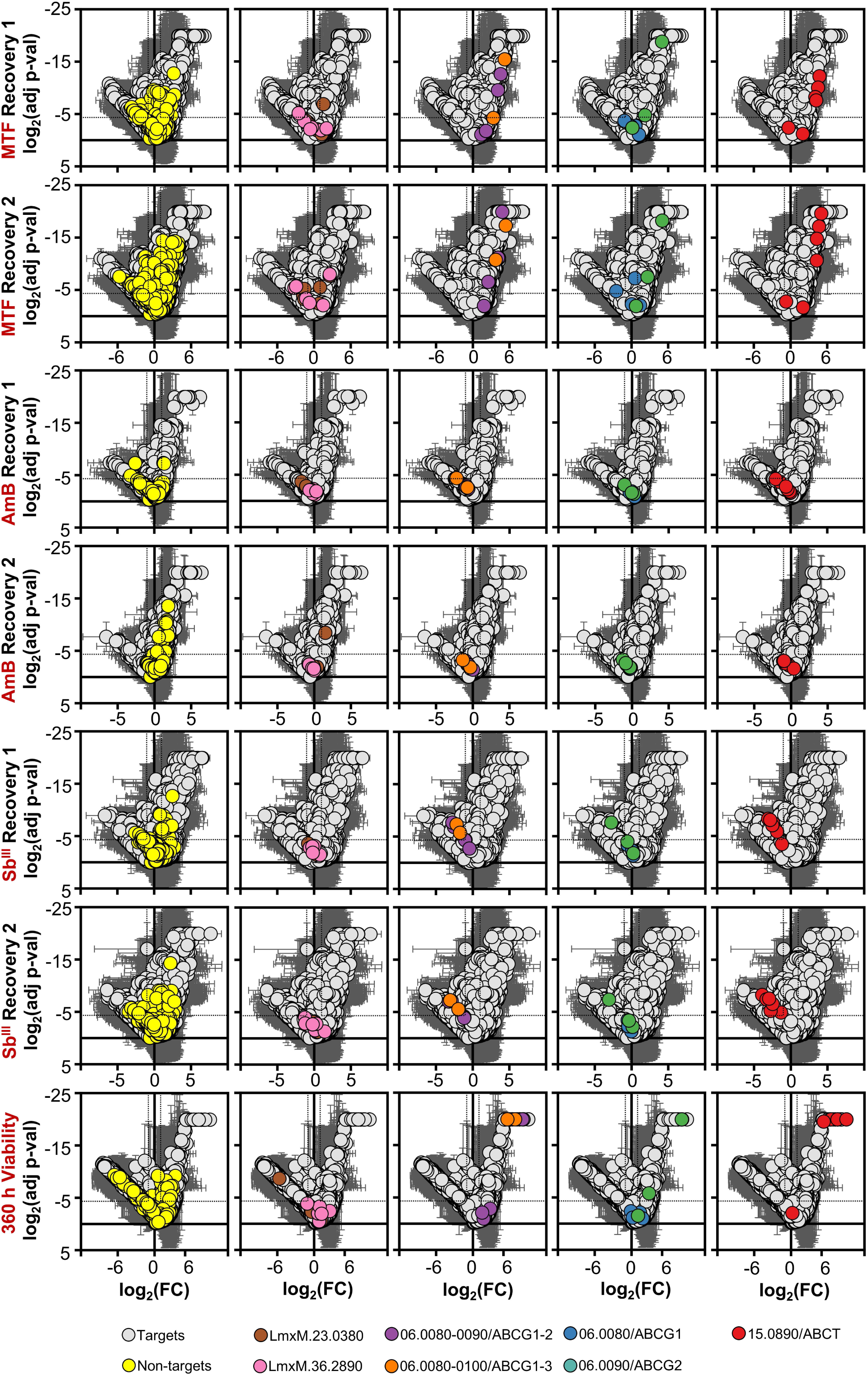
sgRNA distribution of ABCT and ABCG1-3 paralogs highlights collateral sensitivity. Distribution of log_2_ fold change (x-axis) and log_2_ p-values (y-axis) for all 38,422 targeting and 1,000 non-targeting sgRNAs. Error bars represent standard deviation across three biological replicates. Dotted lines mark thresholds for significant enrichment or depletion (log_2_ fold change ≤ -1 or ≥ 1; log_2_ p-value ≤ - 4.3219). Individual sgRNAs targeting specific genes (highlighted below each panel) are annotated. Comparisons shown include, from top to bottom: MTF, AmB, and Sb^III^ recovery fractions vs. their corresponding viability screen time points (192 h vs. recovery 1; 264 h vs. recovery 2), and 0 h plasmid reference vs. the 360 h post-transfection sample.

## SUPPLEMENTARY FILES

**Supplementary File 1. Genome-wide sgRNA library for *L. mexicana*.** File provides details for all CBE sgRNAs used in the genome-wide screen (38,422 targeting and 1,000 non-targeting). Each row corresponds to an individual sgRNA, including its sequence, target gene(s), editing characteristics, and predicted stop codon introduction. If a sgRNA matches multiple geneIDs, information is shown for the first listed target.

**Supplementary File 2. Raw/normalized sgRNA read counts and associated quality metrics.** File contains MAGeCK analysis outputs, including QC summaries, raw sgRNA read counts, and normalized sgRNA counts. QC metrics are plotted in fig. S1E-G and include gini indices, library mapping rate to sequencing data and zero sgRNA counts. R1: recovery 1; R2: recovery 2.

**Supplementary File 3. Individual sgRNA data.** This folder contains MAGeCK-derived data for all 38,422 targeting sgRNAs, along with 1,000 non-targeting control sgRNAs. For each experimental time point, a dedicated folder is provided that includes: (1) MAGeCK Test output files for three biological replicates (replicate vs. control comparison, as indicated in filenames), and (2) a summary file compiling the log_2_ fold change and log_2_ p-value for each replicate. The summary additionally calculates the average and standard deviation for each sgRNA. This dataset serves as the basis for all individual sgRNA plots shown in the study and is available upon individual geneID search on https://www.LeishBASEeditDB.net. Rec1: recovery 1; Rec2: recovery 2.

**Supplementary File 4. Median sgRNA set data.** This folder contains MAGeCK-derived data for the median values of sgRNA sets targeting 7,718 unique or multi-gene targets, along with 200 non-targeting control sgRNA sets. For each experimental time point, a dedicated folder is provided that includes: (1) MAGeCK Test output files for three biological replicates (replicate vs. control comparison, as indicated in filenames), and (2) a summary file compiling the log_2_ fold change and log_2_ p-value for each replicate. The summary additionally calculates the average and standard deviation for each sgRNA set. This dataset serves as the basis for all median sgRNA set plots shown in the study and represents the core content uploaded to https://www.LeishBASEeditDB.net. Rec1: recovery 1; Rec2: recovery 2.

**Supplementary File 5. PW and MLE fitness rank analysis of median sgRNA set data.** For fitness rank analysis, the log_2_ fold changes from the 360 h versus 0 h plasmid comparison (**Sheet 5A**) or MLE z-scores (**Sheet 5B**) were ranked from highest to lowest. Depleted and enriched genes, corresponding to fitness loss and gain, respectively, were defined based on relative enrichment compared to the non-targeting control distribution (outside the 95% population interval of the non-targets). Columns highlighted in red correspond to those plotted in fig. 1D (PW; **Sheet 5A**) and 1F (MLE; **Sheet 5B**).

**Supplementary File 6. Promastigote fitness GO term ranking.** This file contains two sheets describing GO term class analyses.

**Sheet 6A - PRO GO term fitness effects:** Column 1: 2,243 unique Gene Ontology (GO) term classes identified for *L. mexicana*. Column 2: Corresponding GO IDs. Column 3: Total number of genes annotated per GO term. Column 4: Number of non-screened genes per GO term. Column 5: Number of screened genes per GO term. Column 6: Number of multi-gene-targeting sgRNA sets per class. Column 7: Number of genes not associate to a single fitness effect. Columns 8-13: Number and percentage of genes classified as depleted, unchanged, or enriched for each GO term.

**Sheet 6B - *L. mexicana* GO Term annotations:** List of 8,145 annotated genes in *L. mexicana*, including predicted protein products and associated GO term annotations (TriTrypDB v59) (*135, 136*).

**Supplementary File 7. Combined drug recovery median sgRNA set data.** Combined scores (log_2_ fold change of recovery 1 and 2 averaged) and corresponding median sgRNA set ranks for each drug.

**Supplementary File 8. Sterol profiling.** This file provides sterol profiling data for ΔABRM7 (LmxM.25.2375), ΔABRM8 (LmxM.34.0270), and parental *L. mexicana* (PARCas12a) cell lines.

**Sheet 8A - RAW peak area:** Contains raw sterol abundance data.

**Sheet 8B - Normalized peak area:** Shows the same sterol data from Sheet 6A normalized to total sterol content, enabling relative comparisons across samples.

**Sheet 8C - Selected sterol profile:** Includes a focused set of six key sterol species. Data are shown for each biological replicate, along with calculated averages, standard deviations, and results of two-tailed t-tests. Where no signal was detected, a pseudo-value of 0.1 was used for statistical analysis.

**Supplementary File 9. Lipidomics profiling.** This file contains two sheets summarizing the lipidomic alterations in selected *L. mexicana* knockout strains compared to the parental line (PARCas12a).

**Sheet S9A - Lipidomics data:** Contains raw and processed lipidomic measurements for parental cells and null mutants of ADK (LmxM.04.0960), PRM11 (LmxM.33.3955), SPT-like (LmxM.33.3740), and DHCS (LmxM.30.1780). Relative intensities for detected lipid isotopes are shown across four biological replicates for each strain. Isotopes are grouped by lipid class, ionization mode, and include additional metadata such as adducts, molecular formula, mass error, and isotope similarity. For each isotope, average intensity, standard deviation, and Mann-Whitney U test p-values (comparing mutant vs. parental) are provided. These data correspond to fig. 7E.

**Sheet S9B - Stouffer’s statistics:** Provides combined p-values per lipid class for each mutant, calculated using Stouffer’s method to integrate the individual Mann-Whitney U test p-values of all isotopes within the same class. Lipid classes with p < 0.05 are considered significantly altered and marked accordingly.

**Supplementary File 10. Primer list.** This table contains the full set of primers used to generate and validate the mutant cell lines described in this study. Each entry includes an internal reference code, the corresponding *L. mexicana* gene ID, the annotated gene product, and the gene name used throughout the manuscript.

**Supplementary File 11. Individual drug response data.** This file contains three sheets detailing drug response metrics for each tested mutant strain.

**Sheet S11A - EC_50_:** Lists average EC_50_ values for each mutant, determined by MTT assays, with standard deviations from up to nine replicates. Includes comparisons between mutant strains (knockouts and addbacks) and their controls, as well as addbacks versus their corresponding knockout parent strains.

**Sheet S11B – AQP1:** Includes EC_50_ comparisons between *L. mexicana* and *L. major* AQP1 knockout and addback lines, along with their respective parental strains.

**Sheet S11C - Treatment & Recovery:** Reports the average parasite concentration at the end of the treatment phase (Columns F–K) and the recovery phase (Columns L–Q).

Statistical comparisons between mutants and the parental strain were performed using one-way ANOVA followed by Dunnett’s post hoc test. Comparisons between knockout lines and their respective addbacks were analyzed using a two-tailed t-test. Statistically significant differences are highlighted in green; non-significant results are marked in pink.

